# Systemic neoantigen-specific T cells reveal central determinants of PD-(L)1 blockade efficacy

**DOI:** 10.64898/2026.03.10.710196

**Authors:** Célia Ramade, Noémie Thébault, Clara-Maria Scarlata, Daniel Oreper, Françoise Lauzéral-Vizcaino, Suchit Jhunjhunwala, Bastien Cabarrou, Milena Hornburg, Caroline Fournier, Anna Salvioni, Marie Michelas, Victor Sarradin, Giulia Costanza Leonardi, Virginie Feliu, Myriam Maixent, Lise Scandella, Meng Xiao He, Martine Darwish, Amy Heidersbach, Catherine Ross, Hang Xu, Fanny Bouquet, Catia Fonseca, Tom Lesluyes, Nicolas Congy-Jolivet, Carlos Gomez-Roca, Alejandra Martinez, Christel Devaud, Thomas Filleron, Jean-Pierre Delord, Julien Mazières, Lélia Delamarre, Maha Ayyoub

## Abstract

The contribution of neoantigen-specific T cells to PD-(L)1 efficacy has largely been inferred from tumor mutational burden. We functionally profiled circulating T cell responses against 7,038 predicted HLA-I–restricted and 21,453 HLA-II–restricted neopeptides in 27 patients with advanced non-small cell lung cancer treated with anti–PD-(L)1. CD4 responses were frequent and correlated with neoantigen availability but not clinical benefit. In contrast, the magnitude and breadth of neoantigen-specific CD8 T cell responses were associated with clinical benefit, progression-free and overall survival, independently of tumor mutational burden. Patients mounting coordinated CD4 and CD8 responses experienced improved progression-free survival. Tumors from CD8 responders displayed immune signatures indicative of both T cell priming and effector functions. Circulating neoantigen-specific CD8 T cells recognized endogenously processed antigens, trafficked to tumors, and selectively expanded under therapy while retaining CD28, CD226, and CXCR3 expression. These findings identify coordinated, functionally engaged neoantigen-specific T cell responses as central determinants of PD-(L)1 efficacy.

## Introduction

Immune checkpoint blockade (ICB) targeting the PD-1/PD-L1 axis has become a cornerstone of primary and maintenance therapy across multiple tumor types, and is increasingly used in both adjuvant and neoadjuvant treatment settings^1–5^. In non-small cell lung cancer (NSCLC), this approach is particularly relevant for patients whose tumors lack targetable oncogenic drivers, representing the majority of patients. Despite this major therapeutic advance, a substantial proportion of patients fail to derive durable benefit. From the earliest clinical trials, correlative studies have aimed to elucidate the mechanisms underlying therapeutic efficacy, not only to identify predictive biomarkers but, more critically, to inform the development of rational combination strategies that build upon the capacity of anti–PD-(L)1 therapies to mobilize potent antitumor immune responses. Among the earliest features associated with clinical benefit, intratumoral CD8 T cell infiltration^6^ and high tumor mutational burden (TMB)^1,7^ pointed to neoantigens as potential key targets of the antitumor T cell response.

Neoantigen-specific CD8 and CD4 T cells have been identified as components of spontaneous antitumor immune responses in malignancies with both high^8,9^ and low^10^ TMB. The relevance of these T cells has been further underscored by their association with clinical responses following adoptive transfer of tumor-infiltrating lymphocytes (TILs)^11^ and has motivated the development of T cell therapies targeting neoantigens^12^. In the context of ICB, the role of neoantigen-specific T cells has been inferred not only from the association between TMB and predicted tumor neoantigen burden (TNB) and clinical benefit^1,13–15^, but also from evidence of clonal selection against somatic mutations and neoepitopes during therapy^16–18^. Nevertheless, several studies have questioned the robustness of TMB as a standalone predictor of ICB efficacy^19,20^, and others have emphasized the importance of integrating TMB with features indicative of an ongoing in situ immune response^21–23^. These insights have motivated efforts to directly assess T cell responses to predicted neoantigens. Although such studies have been conducted in limited cohorts of patients, they have supported the clinical relevance of neoantigen-specific T cells as mediators of ICB benefit^16,24,25^. Yet, a comprehensive evaluation of both CD4 and CD8 T cell responses across a broad set of predicted neoantigens, of their functionality and of their association with clinical outcome, remains lacking.

We and others have shown that response to anti–PD-(L)1 relies on the mobilization of two contingents of exhausted T cells, which are themselves direct targets of ICB. Late exhausted CD8 T cells, largely composed of tumor-specific cells residing in the tumor^8,26,27^, can be functionally reinvigorated by PD-(L)1 blockade^26–29^. In contrast, early exhausted tumor-specific CD8 T cells, found both within tumors and in the circulation, possess proliferative capacity upon PD-(L)1 blockade^26,28,30^, a process supported by the restored helper function of exhausted CD4 T cells^31^, ultimately contributing to the replenishment of T cells at the tumor site. Circulating tumor-specific T cells are not only direct targets of anti–PD-(L)1 therapy and contributors to clinical responses, they also mirror the antitumor response occurring at the tumor site, as shared clonotypes are frequently detected across both compartments^25,32^. Owing to their accessibility, circulating T cells offer a powerful and scalable window into the tumor-specific T cell repertoire, enabling comprehensive analysis in large patient cohorts, particularly in advanced or metastatic settings, where viable TILs are rarely available during systemic anti–PD-(L)1 therapy.

Here, in a cohort of 27 patients with locally advanced or metastatic NSCLC treated with anti–PD-(L)1, we performed a comprehensive functional assessment of circulating neoantigen-specific T cells. We interrogated CD4 T cell responses using 2,103 long peptides encompassing 21,453 predicted HLA-II binders and CD8 T cell responses against 7,038 predicted HLA-I–restricted neopeptides, collectively covering 60.5% and 78.3% of all mutations predicted to generate HLA-II and HLA-I binders, respectively. Neoantigen-specific CD4 T cell responses were detected in a large proportion of patients and were associated with neoantigen availability. In contrast, circulating neoantigen-specific CD8 T cells were associated with clinical benefit, progression-free survival (PFS), and overall survival (OS), independently of TMB and predicted TNB. Patients with detectable circulating CD8 T cell responses differed from those without responses in the intrinsic features of their top predicted HLA-I neopeptides and in tumor immune transcriptomic signatures associated with antigen processing and interferon signaling. Circulating neoantigen-specific CD8 T cells recognized endogenously processed antigens, were detectable within tumors, and exhibited late memory differentiation, effector function, and an intermediate exhausted phenotype. Finally, longitudinal ex vivo tracking using peptide–HLA-I (pHLA-I) tetramers demonstrated selective expansion of neoantigen-specific CD8 T cells during therapy in clinical responders, with a predominance of T effector memory (T_EM_; CCR7^-^CD45RA^-^) relative to T_EMRA_ (CCR7^-^CD45RA^+^) phenotype, associated with retention of CD28 and CD226 and expression of CXCR3.

## Results

### Patient cohort and characterization of the tumor mutational burden and predicted neoantigen landscape

Twenty-seven patients with locally advanced or metastatic NSCLC, 25 with lung adenocarcinoma (LUAD) and 2 with lung squamous cell carcinoma (LUSC), treated with anti–PD-(L)1, were prospectively included in the study (Table 1 and Figure 1A). Based on the standard-of-care during the accrual period, most patients (*n* = 22) received single-agent therapy; the remaining 5 patients had an immunotherapy-chemotherapy (carboplatin/pemetrexed or carboplatin/paclitaxel) combination (Table 1). Immunotherapy was the first line treatment for 15 patients, whereas 10 and 2 patients received one and two prior lines of therapy, respectively (Table 1). The clinical benefit (CB) group (*n* = 12) comprised 11 patients who achieved a best objective response (BOR) of complete response (CR, *n* = 1) or partial response (PR, *n* = 10) according to RECIST v1.1 criteria, as well as one patient with stable disease (SD) and progression-free survival (PFS) ≥ 6 months (Figure 1A and 1B). The no clinical benefit (NCB) group (*n* = 15) included 12 patients with progressive disease (PD) and 3 patients with SD and PFS < 6 months (Figure 1A and 1B).

**Figure 1.**
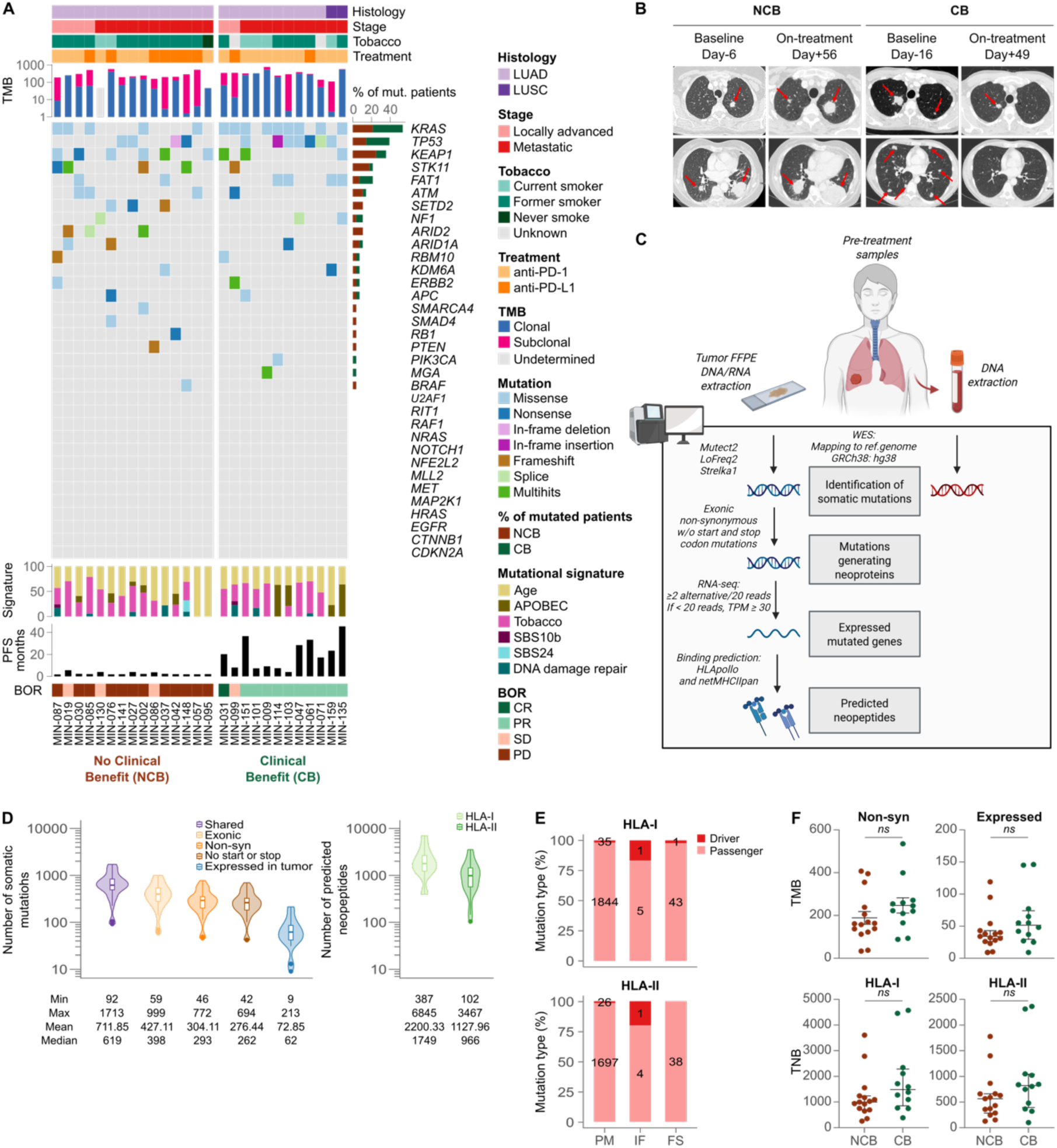
Patient cohort and tumor mutational and neoantigen landscape. (A) Clinical characteristics and tumor genomic features (*n* = 27). (B) Representative computed tomography (CT) scans from one patient with progressive disease (PD) and no clinical benefit (NCB), and one patient with partial response (PR) and clinical benefit (CB), shown at baseline and on treatment. Red arrows indicate tumor lesions. (C) Schematic overview of the neoantigen prediction pipeline. (D) Left: number of tumor somatic mutations after sequential filtering steps (predicted by ≥ 2 variant calling software (Shared), exonic, non-synonymous (Non-syn), not resulting in loss of the initiation codon or gain of a stop codon (No start or stop), and detected in tumor RNA-Seq data (Expressed)). Right: number of HLA-I– and HLA-II–restricted neopeptides (9–11 and 9–15 amino acids, respectively) derived from expressed mutations. Minimum (Min), maximum (Max), mean, and median values are shown. (E) Distribution of mutation type (point mutations [PM], in-frame indels [IF], and frameshift indels [FS]) as well as proportions of driver vs passenger mutations leading to predicted HLA-I and HLA-II binders. (F) Normalized tumor mutational burden (TMB, non-synonymous and expressed) and tumor neoantigen burden (TNB, HLA-I and HLA-II) stratified by clinical benefit (NCB, *n* = 15, red; CB, *n* = 12, green). Mann–Whitney test (median and IQR), except for non-synonymous TMB (unpaired t-test, mean ± SEM). *ns*, not significant.

**Table 1.**
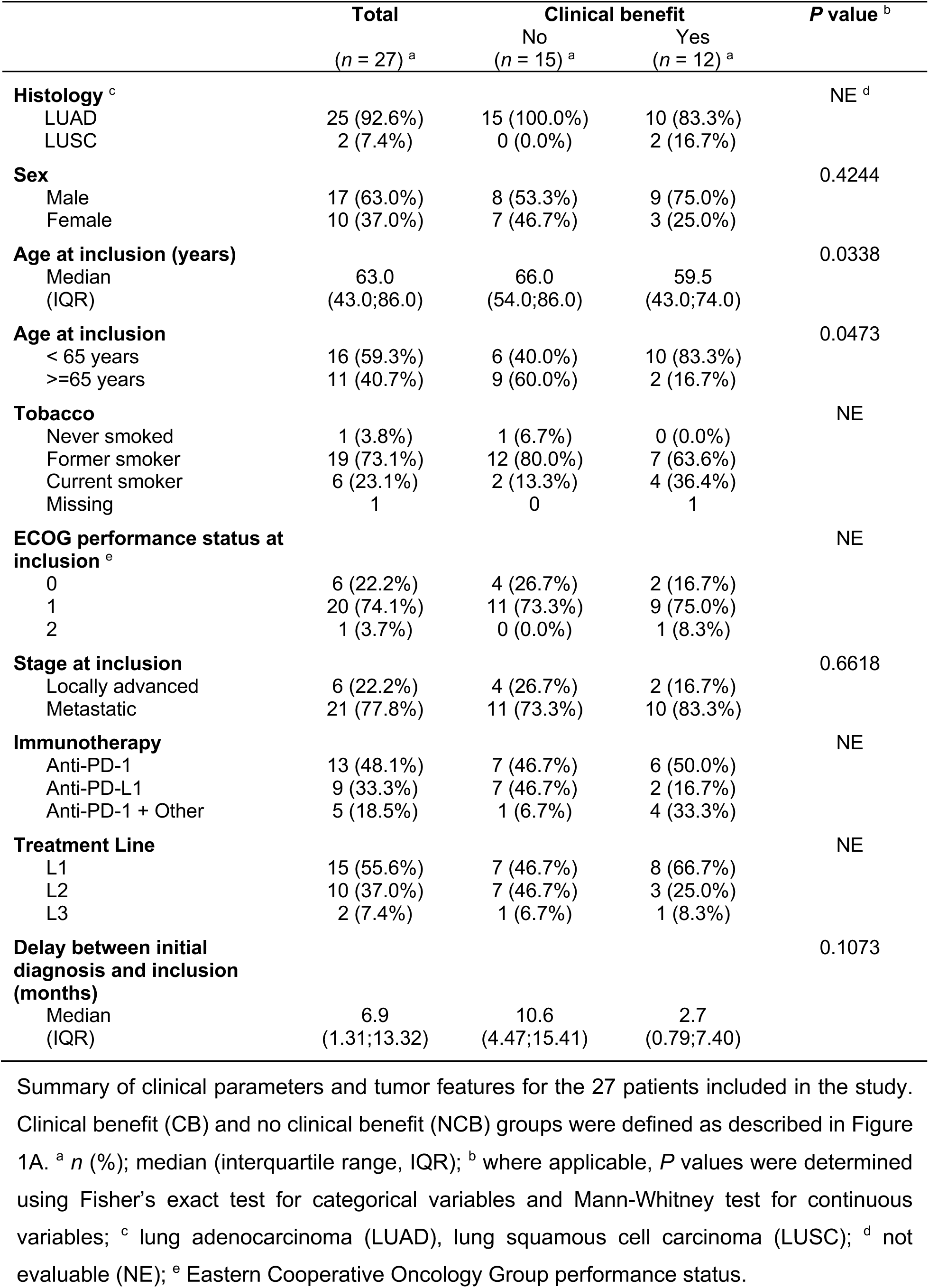
Clinical and tumor characteristics of the patient cohort.

Whole-exome sequencing (WES) was performed on pre-treatment tumor DNA (tumor exome) and matched peripheral blood mononuclear cell (PBMC) DNA (germline exome) (Figure 1C). Tumor mutational burden (TMB) was determined through variant calling using Mutect, LoFreq, and Strelka (Figure 1C). Exonic non-synonymous mutations, detected by at least two of the three algorithms^33^, and that did not result in loss of the initiation codon or gain of a stop codon were retained for downstream analyses (Figure 1C and 1D). The median number of exonic non-synonymous mutations was consistent with those reported in previous series^13,33^. To identify mutations expressed in the tumor and thus potentially generating neoproteins, we performed RNA sequencing (RNA-seq) on pre-treatment tumor RNA (Figure 1C). Mutations were considered expressed if the mutated locus was covered by ≥ 20 RNA reads and included ≥ 2 variant reads. For positions with < 20 reads, expression at the gene level was considered using a threshold of transcripts per million (TPM) ≥ 30^34^. Sequences harboring selected mutations were transcribed in silico for subsequent neopeptide prediction.

HLA-I and -II alleles for each patient were inferred from tumor and germline WES data using HLA-HD^35^. In cases of ambiguity, conventional HLA typing was performed on PBMC-derived DNA. For each patient, HLA-I presentation scores of 9–11-mer neopeptides were predicted using HLApollo^36^, which integrates parameters beyond binding affinity to estimate antigen presentation. In parallel, binding of 9–15-mer neopeptides to HLA-II alleles was predicted using netMHCIIpan^37^. The median numbers of predicted HLA-I and HLA-II neopeptides were high, at 1,749 and 966 per patient, respectively (Figure 1D). Notably, some neopeptides were predicted to bind multiple HLA alleles in the same patient, resulting in a higher number of distinct neoepitopes, defined as unique neopeptide/HLA combinations, than the unique neopeptides presented in Figure 1D. Most predicted neopeptides were derived from point mutations (PMs) rather than in-frame indels (IFs) or frameshift indels (FSs), and from passenger rather than driver mutations (Figure 1E).

To evaluate the association between clinical benefit and TMB or predicted TNB in a manner accounting for read depth heterogeneity, mutations were considered only if located in regions with a coverage of ≥ 50 reads in all 27 tumor WES data. No significant association was observed between TMB and response to therapy, regardless of whether RNA expression of the mutations was taken into account (Figure 1F and Table S1). Similarly, no association was found between predicted HLA-I or HLA-II TNB and clinical response (Figure 1F and Table S1). Whereas these results, which were in contrast with published reports^38^, may reflect the small size of the cohort, they warranted further investigation of T cell responses to the predicted neoepitopes to explore whether responsiveness to anti–PD-(L)1 was more robustly associated to neoantigen-specific T cell responses.

### Feasibility and clinical relevance of a comprehensive assessment of circulating neoantigen-specific T cell responses

To comprehensively assess CD4 T cell responses to neoantigens, we considered up to 50 mutations per patient. These were selected according to the binding score of the best neopeptide-HLA-II pair among all predicted HLA-II binders encoded by each mutation (netMHCIIpan). Given that in vitro recall of CD4 T cell responses is permissive to peptide lengths exceeding the core HLA-II binding sequence, we designed extended peptide sequences encompassing each mutated residue (PMs and IFs) and all overlapping predicted HLA-II binders containing the mutation (Figure S1A and S1B). According to their length, extended sequences were synthesized as one peptide or as overlapping 20-mers (Figure S1A and S1B). Peptide sequences resulting from FSs were entirely covered by 20-mers overlapping by at least 10 amino acids (Figure S1C). For each patient, the ensuing long peptides were synthesized and pooled. The long peptide pool was used to stimulate CD4 T cells, isolated from pre- and on-treatment PBMCs, in the presence of autologous CD14 monocytes as antigen-presenting cells (APCs) (Figure 2A). After 11 days of culture, cells were re-stimulated or not with the same peptide pool, and antigen-specific T cells were identified by intracellular cytokine staining (Figure 2A and 2B). CD4 T cell responses were readily detectable in 18 of 27 patients (66.7%), including 9 of 12 patients with clinical benefit (CB) and 9 of 15 with no clinical benefit (NCB) (Fisher’s exact test, *P* = 0.68) (Table S1), with no significant difference in response magnitude between the two groups (Figure 2C and Table S1).

**Figure 2.**
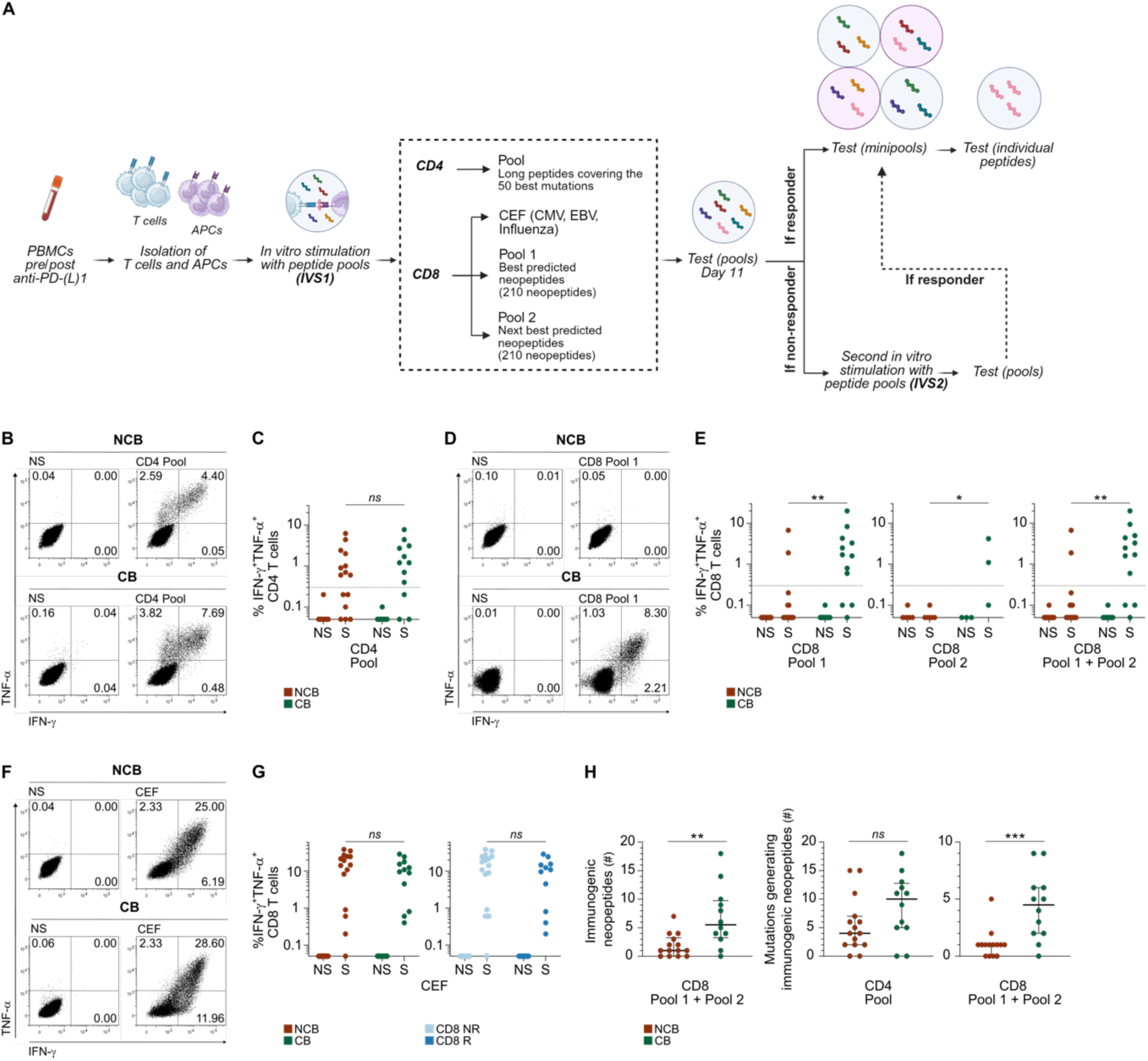
Feasibility and clinical relevance of a comprehensive assessment of circulating neoantigen-specific T cell responses. (A) Schematic overview of the in vitro stimulation (IVS) assay used to assess the magnitude and breadth of CD4 and CD8 T cell responses. Cultures of patient-derived PBMCs were stimulated with pools of predicted neopeptides or individual neopeptides and tested by intracellular cytokine staining and flow cytometry. CEF, CMV, EBV, and influenza peptides. (B) Representative dot plots of cytokine production (IFN-γ and TNF-α) in CD4 T cells from day 11 IVS1 cultures under unstimulated (NS) and stimulated (S) conditions, shown for one non-clinical benefit (NCB) and one clinical benefit (CB) patient. Numbers indicate the percentage of cells in each quadrant. (C) Proportions of IFN-γ^+^TNF-α⁺ CD4 T cells in day 11 IVS1 cultures stratified by clinical benefit. Patients were classified as CD4 responders if the proportion of cytokine⁺ cells in the S condition was ≥ 0.3% (dotted line). (D) Representative dot plots of CD8 T cell responses to pool 1 from one NCB and one CB patient. (E) Proportions of IFN-γ⁺TNF-α⁺ CD8 T cells in day 11 IVS1 cultures for pool 1 (left, *n* = 27), pool 2 (center, *n* = 8), and the sum of responses to both pools (right, *n* = 27), stratified by clinical benefit. Patients were classified as CD8 responders if the proportion of cytokine⁺ cells in the S condition was ≥ 0.3% (dotted lines). (F) Representative dot plots of CD8 T cell responses to CEF from one NCB and one CB patient. (G) Proportions of IFN-γ⁺TNF-α⁺ CD8 T cells in day 11 CEF cultures (*n* = 27), stratified by clinical benefit and CD8 T cell response to neopeptides (as determined in E, right). (H) Number of immunogenic CD8 neopeptides per patient (left), and number of tumor mutations giving rise to immunogenic CD4 (center) or CD8 (right) neopeptides. Median and IQR (H). Mann–Whitney test (S conditions (C,E,G) and H). \**P* < 0.05, \*\**P* < 0.01, \*\*\**P* < 0.001; *ns*, not significant

To assess CD8 T cell responses, we selected the top 210 neopeptides with the highest predicted HLA-I presentation scores (HLApollo) for each patient and, for 8 patients (3 CB and 5 NCB), an additional set of the next-best 210 neopeptides was also included (Figure 1D). These synthesized peptides, corresponding to the exact length of predicted HLA-I binders, were pooled for each patients and used to stimulate circulating CD8 T cells according to a protocol analogous to that applied for CD4 T cells (Figure 2A). In cultures stimulated with the top 210 neopeptides (Pool 1), CD8 T cell responses, measured by IFN-γ and TNF-α production (Figure 2D), were observed in 11 of 27 patients (40.7%), including 9 of 12 CB and only 2 of 15 NCB patients (Fisher’s exact test, *P* = 0.002) (Table S1). The second neopeptide set (Pool 2) failed to induce an observable response in any of the 5 NCB patients, who did not respond to Pool 1, whereas Pool 2 did reveal additional neopeptide-specific responses in two of the three tested CB patients, who already exhibited responses to Pool 1 (Figure 2E). Taken together, these results indicated that the top 210 neopeptides, representing on average ∼10% of all predicted HLA-I binders, provided a representative measure of CD8 T cell reactivity to tumor neoantigens. The magnitude of CD8 T cell responses to neopeptides was significantly associated with clinical benefit, whether considering responses to Pool 1 alone, Pool 2 alone, or both combined (Figure 2E and Table S1). Notably, the absence of neopeptide-specific T cell responses in NCB patients could not be attributed to impaired immune fitness, as 26 of 27 patients mounted responses to control viral antigens (CMV, EBV, influenza), with comparable response magnitudes between the CB and NCB groups and between CD8 neopeptide responders and non-responders (Figure 2F and 2G).

To assess the breadth of response, that is, the number of distinct neopeptides recognized by CD8 and CD4 T cells, we employed a deconvolution strategy as previously described^39^. Minipools (MPs) containing on average 20 neopeptides, each sharing a single neopeptide with another MP, were generated and used to stimulate day 12 cultures, followed by intracellular cytokine staining (Figure 2A and Figure S2A). Neopeptides common to all positive MPs were then tested individually to confirm reactivity (Figure 2A and Figure S2B). For patients in whom CD8 or CD4 T cell responses were undetectable after the initial in vitro stimulation, a second round of stimulation was performed using the same neopeptide pools in the presence of autologous APCs (Figure 2A). Deconvolution of positive cultures was then carried out using the same strategy. The breadth of CD8 responses, whether quantified by the number of immunogenic neopeptides or by the number of tumor mutations encoding them (since a single mutation can give rise to multiple immunogenic neopeptides), was significantly greater in CB than in NCB patients (Figure 2H). In contrast, the breadth of CD4 T cell responses, quantified by the number of tumor mutations encoding immunogenic long peptides, did not differ between CB and NCB patients (Figure 2H).

The robustness of these observations is underscored by the comprehensiveness of the approach. Across the full patient cohort, CD4 T cell responses were evaluated using 2,103 long peptides encompassing 21,453 predicted HLA-II binders. Tested CD4 sequences were derived from 1,068 mutations, representing 60.5% of all mutations predicted to encode HLA-II binders. CD8 T cell responses were tested against 7,038 neopeptides derived from 1,510 mutations, representing 78.3% of all mutations predicted to encode HLA-I binders. Together, these results establish the feasibility of comprehensively interrogating circulating neoantigen-specific T cell responses and identify CD8 T cell reactivity as a key correlate of clinical benefit, warranting further analysis of its association with patient survival.

### Association of circulating neoantigen-specific T cell responses with progression-free and overall survival under anti–PD-(L)1

Among the clinical parameters assessed for association with clinical benefit (as defined in Figure 1A and 1B), only age at inclusion reached statistical significance, younger patients being more likely to experience benefit (Table 1). The limited number of patients in certain subgroups precluded meaningful analysis of histological subtype, tobacco use, ECOG performance status at inclusion, and of the type of immunotherapy received (Table 1). Although the number of prior treatment lines could not be analyzed across all three categories (Table 1), no difference in clinical benefit was observed between patients receiving anti–PD-(L)1 as first-line therapy and those treated in later lines (Chi-squared test, *P* = 0.299). Additionally, the interval between initial diagnosis and initiation of immunotherapy was not associated with clinical benefit (Table 1). Finally, the mutational signature (age, APOBEC, tobacco) and oncogenic mutations (Figure 1A) were not associated with clinical outcome.

The median PFS and OS in the cohort were 4.0 months (95% CI, 2.0–8.0) and 23.2 months (95% CI, 12.6–38.8), respectively (Figure 3A and 3B). Continuous variable analyses showed no significant association between TMB or TNB and clinical benefit (Figure 1F and Table S1). However, when patients were stratified using the median TMB or TNB values, an admittedly arbitrary threshold, high HLA-I TNB was associated with prolonged PFS, though not with OS (Figure 3C and 3D). In contrast, both the magnitude and breadth of CD8, but not CD4, T cell responses were significantly associated with clinical benefit in continuous analyses (Figure 2C, 2E and 2H and Table S1). To assess the impact of T cell responses on survival, patients were classified as T cell responders or non-responders based on the day 11 in vitro stimulation assay (Figure 2E). CD8 responders experienced significantly longer PFS and, notably, OS, compared to non-responders, whereas CD4 responses were not associated with survival (Figure 3E and 3F). A swimmer plot further illustrates the clinical benefit of CD8 T cell responses to neoantigens, highlighting durable disease control in CD8 responders (Figure 3G). Of note, among the 11 CD8 responders, 10 also exhibited CD4 T cell responses whereas of the 16 CD8 non-responders only 8 exhibited CD4 responses (Chi-squared test, *P* = 0.0267). Double-responder patients displayed longer PFS and a trend toward longer OS (*P* = 0.0545), compared to patients with CD4 responses alone or to those lacking detectable T cell responses (Figure 3E and 3F), supporting a contribution of coordinated CD4 and CD8 T cell responses to clinical outcome.

**Figure 3.**
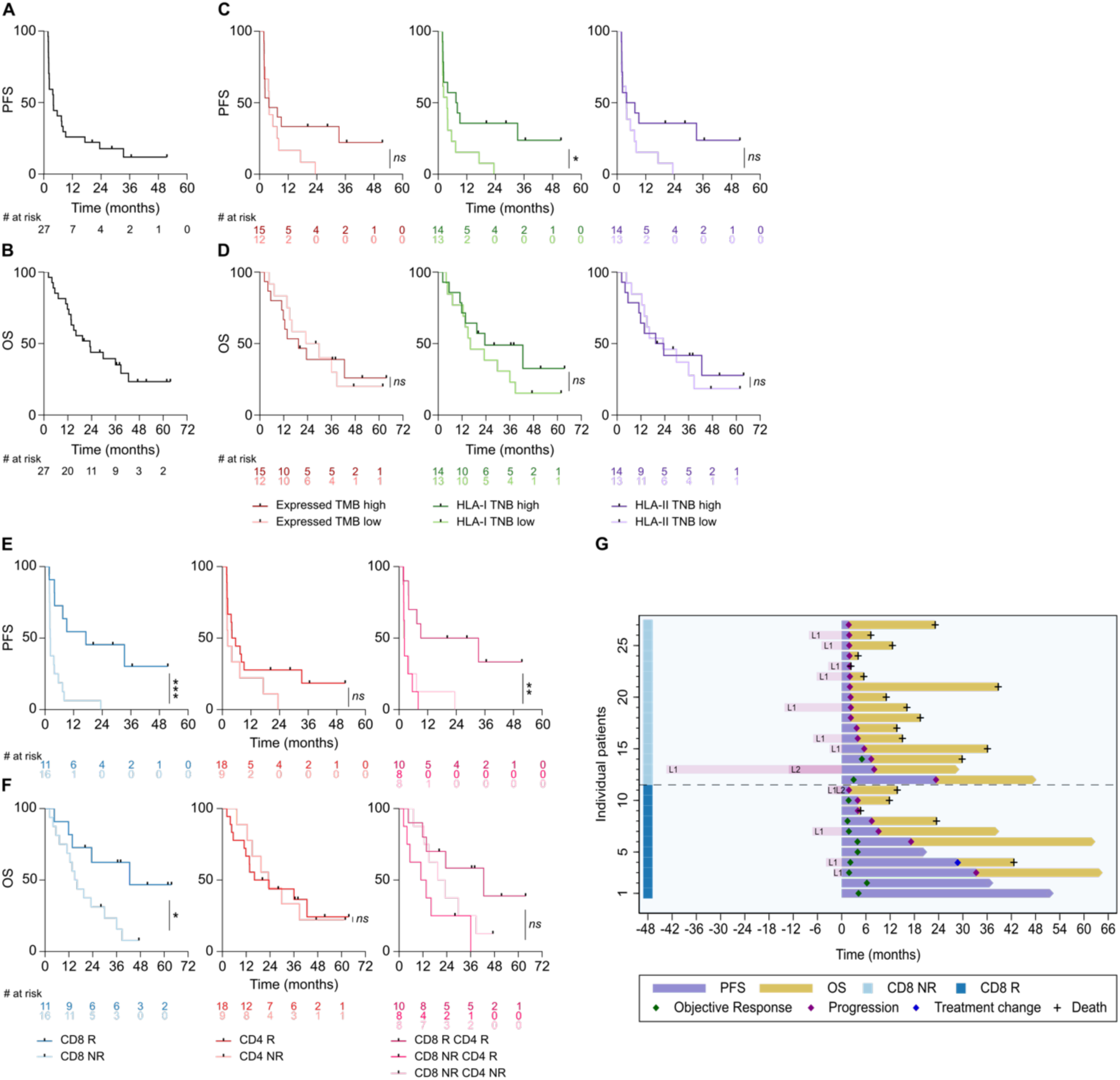
Association of circulating neoantigen-specific T cell responses with progression-free and overall survival under anti–PD-(L)1. (A,B) Progression-free survival (PFS, A) and overall survival (OS, B) (*n* = 27). (C,D) PFS (C) and OS (D) of patients stratified by the median of the normalized expressed tumor mutational burden (TMB, left), HLA-I tumor neoantigen burden (TNB, middle), and HLA-II TNB (right). (E,F) PFS (E) and OS (F) according to circulating CD8 (from Figure 2E) and CD4 (from Figure 2C) T cell responses to neoantigens. (G) Clinical course of all patients, grouped by CD8 T cell responder status (dark blue: responders, *n* = 11; light blue: non-responders, *n* = 16). Log-rank test. \**P* < 0.05, \*\*\**P* < 0.001; *ns*, not significant.

Together, these continuous and survival parameter analyses establish coordinated neoantigen-specific T cell responses as central effectors of immunotherapy efficacy, beyond neoantigen availability alone, and provide the rationale for dissecting the features that underlie their clinical impact.

### Intrinsic neoantigen features and tumor immune context contribute to CD8 T cell priming

Like TMB and TNB-II (Figure 1F and Table S1), CD4 T cell responses, detected in 18 patients, were not associated with response to therapy (Figure 2C and 2H). Both the magnitude and breadth of CD4 responses were positively correlated with TMB and TNB-II (Figure 4A and 4B), consistent with neoantigen availability being a major determinant of CD4 T cell priming. In contrast, the differences in CD8 T cell responses observed between CB and NCB patients (Figure 2E and 2H and Table S1) occurred despite the absence of association between clinical benefit and either TMB or TNB-I (Figure 1F and Table S1). This dissociation was further supported by the lack of correlation between the magnitude or breadth of CD8 responses and TMB or TNB-I, within both clinical and CD8 responder groups (Figure 4C and 4D). Together, these findings indicate that clinical benefit from anti–PD-(L)1 therapy is not explained by neoantigen availability alone, but rather by the effective engagement of an antitumor CD8 T cell response.

**Figure 4.**
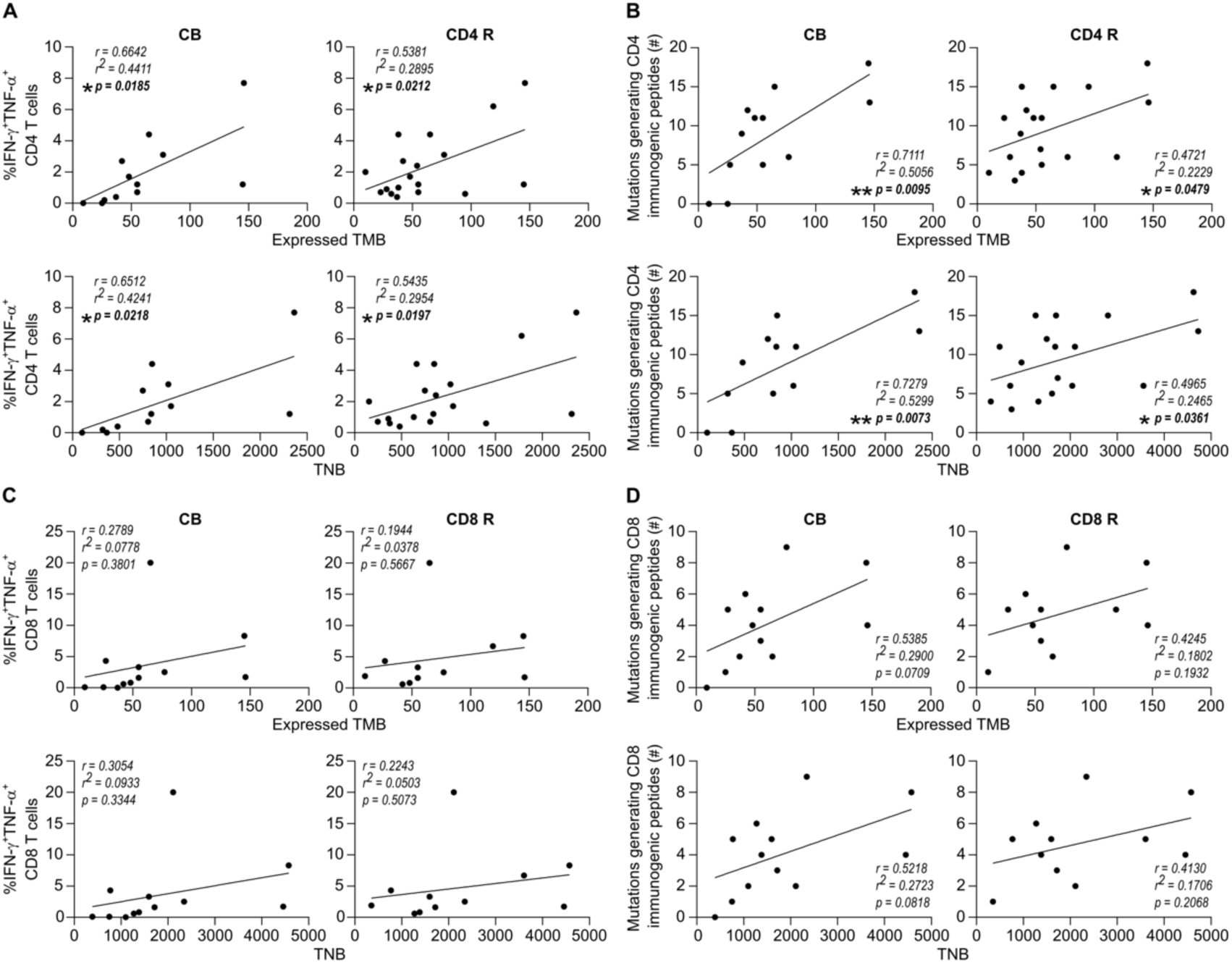
Magnitude and breadth of neoantigen-specific CD8 T cell responses do not correlate with TMB or TNB. Correlations were performed in patients with clinical benefit (CB, *n* = 12; Figure 1A) or with T cell responses to neopeptides (CD4 responders [R], *n* = 18, Figure 2C; CD8 R, *n* = 11; Figure 2E, left). (A,B) Correlation between normalized expressed tumor mutational burden (TMB) or HLA-II tumor neoantigen burden (TNB) and the magnitude (A) and breadth (B) of CD4 T cell responses. (C,D) Correlation between normalized expressed TMB or HLA-I TNB and the magnitude (C) and breadth (D) of CD8 T cell responses to pool 1. Pearson correlation test. \**P* < 0.05, \*\**P* < 0.01

To investigate whether differences in the intrinsic properties of neoantigens could account for CD8 T cell priming, we analyzed all tested neopeptides across the cohort. Among the 7,038 CD8 neopeptides evaluated, derived from 1,510 mutations, 108 (1.5%) were immunogenic. In parallel, the 2,103 long peptides assessed for CD4 reactivity were derived from 1,068 mutations, of which 187 (17.5%) encoded immunogenic neopeptides. As expected, immunogenic CD8 neopeptides exhibited features classically associated with immunogenicity, including improved predicted HLA-I presentation (HLApollo), higher binding affinity (netMHCIpan, mixMHCpred), enhanced processing, increased peptide-HLA complex stability, and higher predicted likelihood of T cell recognition (Figure S3A–S3D). In contrast to previous reports^34^, total gene expression was not associated with immunogenicity (Figure S3E and S3F). Instead, both CD4 and CD8 immunogenic neopeptides preferentially originated from transcripts with higher mutant-to-total RNA ratios (Figure S3E and S3F). Among the 243 mutations encoding immunogenic neopeptides, most elicited either CD4 or CD8 T cell responses exclusively, with limited overlap (Figure S3G). Immunogenic CD4 neopeptides were more frequently derived from clonal mutations, whereas mutation clonality did not influence CD8 responses (Figure S3H). Consistent with prior studies^40^, most immunogenic neopeptides originated from passenger rather than driver mutations and from PMs, in line with their overall predominance among predicted neopeptides (Figure 1E and Figure S3I).

To determine whether patients mounting CD8 T cell responses harbored a qualitatively superior neoantigenic landscape, we compared the binding, presentation, processing and recognition scores, as well as mutant-to-total RNA ratios (Figure S3A–S3E), of the top 210 predicted HLA-I neopeptides (corresponding to Pool 1, Figure 2A) between CD8 responders and non-responders. Across these parameters, predicted neoantigen quality was higher in CD8 responders (Figure S3J–S3N). However, a substantial fraction of neopeptides from non-responders displayed characteristics comparable to those of actually immunogenic neopeptides (Figure S3A–S3E), indicating that although intrinsic neoantigen quality contributes to CD8 T cell priming, it does not fully account for the presence or absence of circulating CD8 T cell responses.

Finally, unsupervised clustering of patients based on scaled normalized expression of immune-related genes (including antigen processing and presentation, IFN signaling, and TCR/BCR signaling pathways) segregated patients with and without detectable circulating neopeptide-specific CD8 T cells (Figure S4A). Tumors from CD8 responders were enriched in gene signatures associated with effector CD8 T cells, cytotoxicity, activated memory CD4 T cells, and IFN-γ signaling (Figure S4B), consistent with circulating neoantigen-specific CD8 T cell responses reflecting an ongoing antitumor immune response at the tumor site associated with an active in situ CD4 response. Tumors from CD8 responders were also enriched in signatures linked to CD8 T cell priming, including the immunoproteasome and type I interferon pathways (Figure S4C), providing a potential tumor-intrinsic context associated with these responses.

Together, these findings suggest that intrinsic neoantigen features and tumor immune context contribute to CD8 T cell priming and hence potentially to clinical outcome. The latter may depend on additional parameters related to the quality of primed T cells and their responsiveness to PD-1/PD-L1 axis blockade.

### Circulating T cells recognizing neopeptides are functionally competent and tumor-relevant

To ascertain the relevance of T cell responses detected in the circulation, we further characterized the neopeptide-specific T cells. CD8 T cell responses to 28 neopeptides assessed in in vitro-stimulated cultures from 5 patients demonstrated their specificity for the mutated peptides and virtually no recognition of the corresponding WT sequences encoded by the germline exome (Figure 5A). From two CD8 responder patients, IFN-γ–guided sorting was used to isolate and clone neopeptide-specific CD8 T cells. Assessment of clones specific for 2 different neopeptides from each patient confirmed the specific recognition of the neopeptides, but not the WT sequences, and showed their ability to secrete granzyme B (Figure 5B). Staining of clones with fluorescent pHLA-I tetramers validated the restricting allele and demonstrated clonal purity (Figure 5C). To test recognition of endogenously processed antigen, COS-7 target cells were transfected with plasmids encoding the relevant HLA-I allele and tandem minigenes (TMGs) expressing 27-mer peptides with the mutated amino acid in position 14. Specific clones efficiently recognized the processed neopeptides, as evidenced by upregulation of 4-1BB (Figure 5D), CD107a surface expression due to degranulation (Figure 5E), and secretion of IFN-γ and granzyme B (Figure 5F).

**Figure 5.**
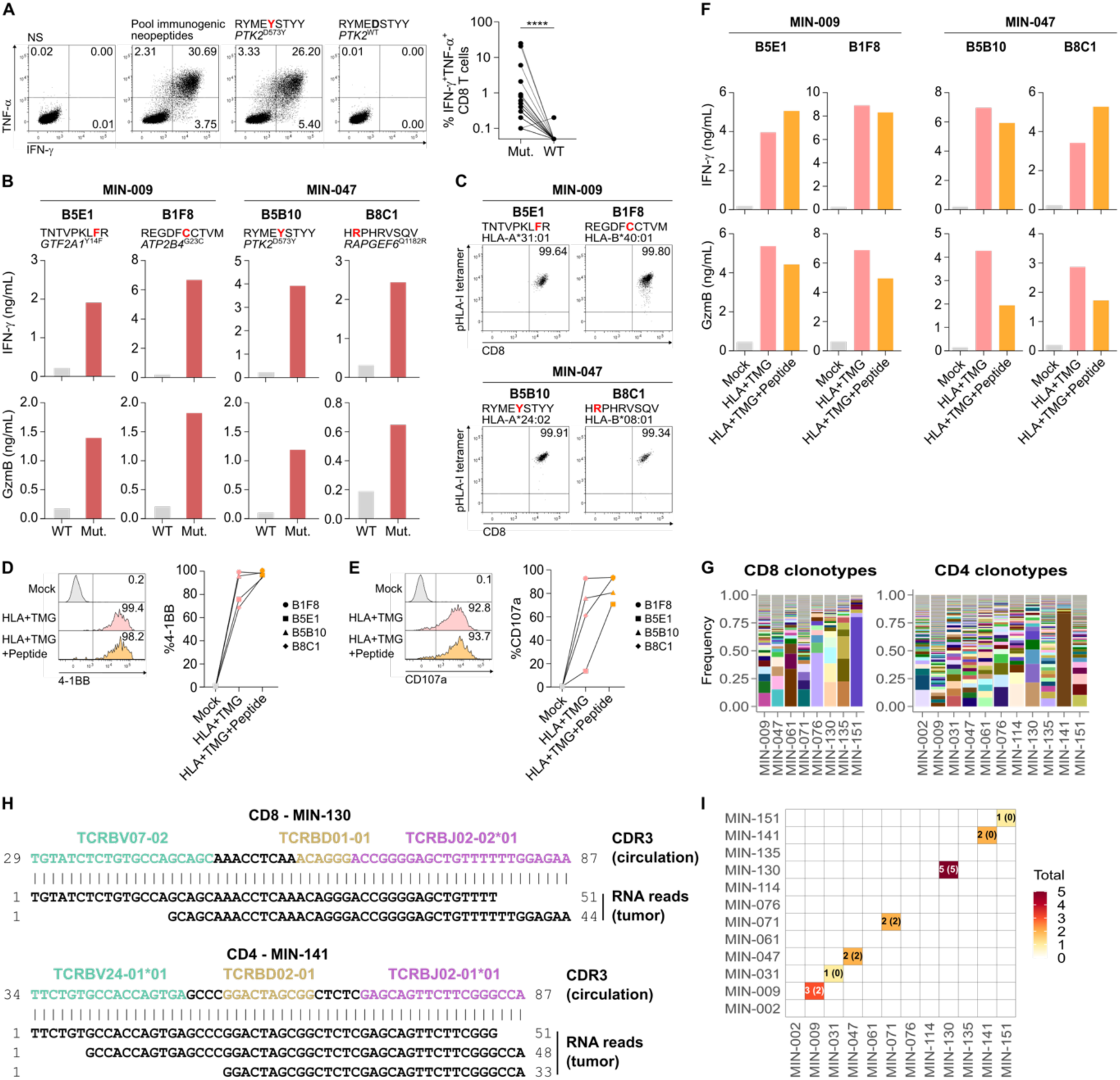
Circulating neopeptide-specific CD8 T cells recognize endogenously processed antigen and localize to the tumor. Patient-derived baseline and on-treatment PBMCs from CD8 and/or CD4 responder patients were stimulated with pools of the individually identified immunogenic neopeptides as in Figure 2A. (A) CD8 T cell cultures were assessed for reactivity to individual mutated neopeptides and their wild-type (WT) counterparts. Left: representative dot plots showing IFN-γ and TNF-α production under unstimulated (NS), pool-stimulated, and individual peptide–stimulated conditions from one patient and specificity. Gene name, mutation position, and peptide sequences are indicated. Right: summary of IFN-γ⁺TNF-α⁺ CD8 T cell responses to 28 matched mutated and WT peptides across 5 patients. (B–F) Neopeptide-reactive CD8 T cells from two patients (MIN-009 and MIN-047) were isolated by IFN-γ–guided sorting, cloned, and functionally characterized. (B) IFN-γ (top) and granzyme B (GzmB, bottom) secretion by T cell clones co-cultured with autologous monocytes pulsed with mutated or WT peptides. (C) Dot plots showing staining of clones with pHLA-I tetramers and anti-CD8. (D–F) Clones were co-cultured with COS-7 cells co-transfected with plasmids encoding the restricting HLA-I allele and tandem minigenes (TMGs) encoding 27-mer peptides centered on mutations, or irrelevant plasmids (Mock). (D,E) Surface expression of 4-1BB (D) and CD107a (E); left, representative histograms; right, summary data. (F) IFN-γ (top) and GzmB (bottom) secretion. Clone name, gene, mutation position, peptide sequence, and HLA restriction (B,C). (G–I) Neopeptide-reactive CD8 and/or CD4 T cells from 12 patients were isolated by IFN-γ–guided sorting and subjected to TCRβ sequencing. (G) Frequencies of clonotypes (TCRβ CDR3 sequence) in sorted CD8 (left) and CD4 (right) populations. (H) Examples of alignment between circulating CDR3 sequences and tumor RNA-seq reads for two patients; V, D, and J segments are shown in green, yellow, and purple, respectively. (I) Correspondence matrix of matched CDR3 reads across patients, colored by total number of circulating clonotypes detected at the tumor site (CD8 CDR3s in parentheses). Wilcoxon matched-pairs rank test (A). \*\*\*\**P* < 0.0001.

IFN-γ–guided sorting was then used to isolate CD8 and CD4 T cells specific for all identified immunogenic neopeptides from 12 patients for whom sufficient material was available (7 with both CD8 and CD4 specific T cells isolated, 1 with CD8 only, and 4 with CD4 only). TCRβ sequencing was performed on sorted specific T cells (Figure 5G). To assess the presence of circulating neopeptide-specific clonotypes at the tumor site, we used BLAST to search each patient’s specific CDR3 sequences in the bulk tumor RNA-seq data from all 12 patients (Figure 5H). Both CD4 and CD8 CDR3s were detectable at the tumor site (Figure 5H and 5I). This approach, although unconventional due to the limited sensitivity of standard RNA-seq for TCR detection, provided robust specificity: neopeptide-specific CDR3s were detected exclusively in the tumor RNA-seq of the matched patient, and not in that of other patients (Figure 5I).

Taken together, these results demonstrate that circulating neoantigen-specific CD8 T cells are bona fide tumor-reactive lymphocytes capable of recognizing endogenously processed antigens and trafficking to the tumor microenvironment, supporting the relevance of their detection in peripheral blood and providing a rationale for their longitudinal ex vivo quantification and phenotypic characterization under therapy.

### Anti–PD-(L)1 therapy promotes expansion of functionally competent neopeptide-specific CD8 T cells in clinical responders

In vitro stimulation experiments enabled the classification of patients as CD8 responders or non-responders (Figure 2E). To avoid missing responses that might have been amplified or lost under therapy, these assays were performed using CD8 T cells obtained both before and during therapy. In addition, CD14 monocytes were used as APCs instead of dendritic cells to minimize in vitro priming of naïve T cells and preferentially recall pre-existing memory responses. Nonetheless, to quantify neopeptide-specific T cells, characterize their phenotype, and monitor their evolution during therapy, ex vivo analyses were required. To this end, we used pHLA-I tetramers to analyze circulating neopeptide-specific CD8 T cells before and on treatment, in patients for whom both appropriate samples and HLA alleles were available. These included three neoepitopes from two NCB patients and eight neoepitopes from six CB patients (Figure 2E).

Tetramer-positive (tetramer^+^) cells were readily detectable ex vivo in all evaluated patients and timepoints (Figure 6A; for gating strategy, see Figure S5A). The frequency of neopeptide-specific CD8 T cells significantly increased during therapy in patients who experienced clinical benefit (Figure 6A). The lower number of specificities assessed ex vivo in the NCB group reflects the absence or limited breadth of CD8 T cell responses in these patients (Figure 2E and 2H). Across all patients and timepoints, tetramer^+^ cells were at late differentiation stages, varying among patients and specificities, with some displaying an effector memory (T_EM_; CCR7^-^CD45RA^-^) phenotype, and others expressing CD45RA, consistent with a T_EMRA_ (CCR7^-^CD45RA^+^) phenotype, whereas cells with a central memory (T_CM_; CCR7^+^CD45RA^-^) phenotype were virtually undetectable (Figure 6B). T_EMRA_ tetramer^+^ cells were overrepresented in NCB patients and the T_EM_/T_EMRA_ ratio was higher in CB patients (Figure 6B), suggesting that enrichment in the T_EMRA_ phenotype was associated with reduced clinical responsiveness to anti–PD-(L)1. Of note, in both patient groups the phenotype of specific T cells did not change under therapy (Figure S5B).

**Figure 6.**
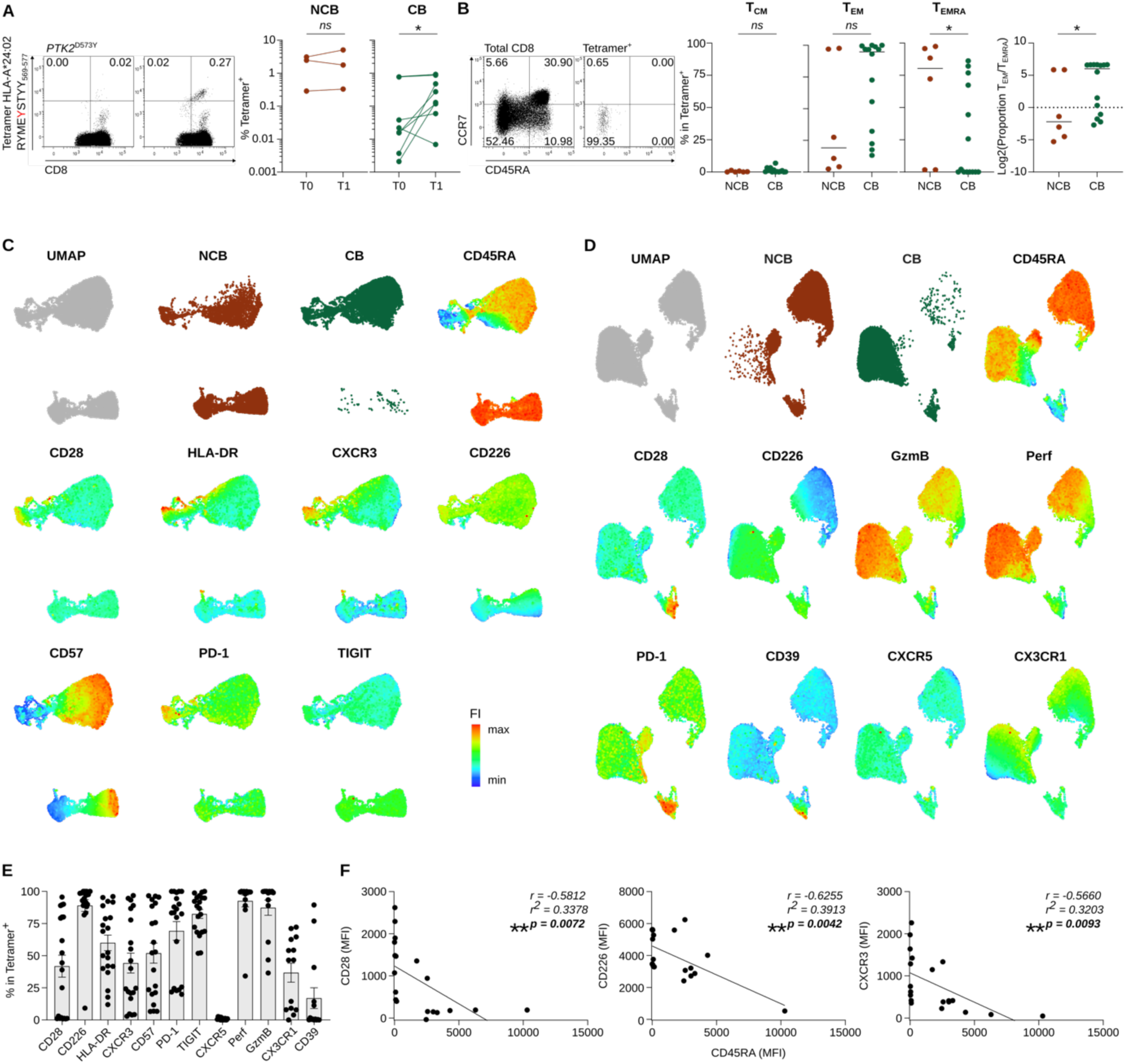
Ex vivo tetramer staining reveals expansion of functional neopeptide-specific CD8 T cells in clinical responders. Circulating CD8 T cells from baseline (T0) and on-treatment (T1) samples were stained ex vivo with personalized pHLA-I tetramers and phenotyped. (A) Left, representative dot plots from baseline (left) and on-treatment (right) samples of one patient with clinical benefit (CB). Gene name, mutation position, neopeptide sequence, and restricting HLA-I allele. Numbers denote the percentage of cells in each quadrant. Right, summary of the frequency of tetramer⁺ cells among memory CD8 T cells (gating in Figure S5A) for 2 patients without clinical benefit (NCB; 3 specificities) and 6 CB patients (8 specificities). (B) Left, dot plots showing CCR7 and CD45RA expression in total CD8 T cells (left) and tetramer⁺ cells (right) from one CB patient. Right, summary of memory tetramer⁺ cells distributed across central memory (T_CM_: CCR7⁺CD45RA⁻), effector memory (T_EM_: CCR7⁻CD45RA⁻), and T_EMRA_ (CCR7⁻CD45RA⁺) subsets. (C,D) Uniform Manifold Approximation and Projection (UMAP) of memory tetramer⁺ CD8 T cells from all specificities and timepoints. Data in C and D correspond to two distinct staining panels (Figure S5C and S5D). Colors indicate clinical group (CB or NCB) or fluorescence intensity (FI) of the indicated markers. (E) Proportions of cells expressing the indicated markers among memory tetramer^+^ cells for all specificities and timepoints (mean ± SEM). (F) Pearson correlations between CD45RA MFI and the MFI of the indicated markers in tetramer⁺ cells of all specificities and timepoints. Perf, perforin; GzmB, granzyme B. Paired t-test (NCB, A), Wilcoxon matched-pairs rank test (CB, A), Mann–Whitney test (B) and Pearson correlation test (F). \**P* < 0.05, \*\**P* < 0.01; *ns*, not significant.

Neopeptide-specific CD8 T cells were further phenotypically characterized (Figure S5C and S5D). Unsupervised clustering of tetramer^+^ cells across all patients and timepoints partially segregated cells from clinical responders and non-responders, though some clusters were shared between groups (Figure 6C and 6D). Tetramer^+^ cells consistently showed high expression of granzyme B and perforin, in line with their differentiation stage and suggesting preserved cytotoxic potential (Figure 6D and 6E). PD-1 and TIGIT were expressed in tetramer^+^ cells from all patients, while CD39 was absent in most samples, consistent with an intermediate exhausted phenotype (Figure 6C–6E) comforted by the relatively higher expression of CX3CR1 compared to CXCR5^26,41–43^ (Figure 6D and 6E). Tetramer^+^ T cells were activated in vivo as evidenced by HLA-DR expression (Figure 6C and 6E). These analyses further confirmed the higher expression of CD45RA in tetramer^+^ cells from NCB patients (Figure 6C and 6D), aligned with their T_EMRA_ phenotype (Figure 6B).

To examine the relationship between the advancement in the differentiation stage and phenotype, we assessed correlations between CD45RA and markers associated with T cell proliferation, namely CD28 and CD226, and CXCR3 for tumor homing, which were all expressed in sizable fractions of tetramer^+^ cells (Figure 6E). CD45RA expression was negatively correlated with CXCR3 (Figure 6F), suggesting reduced tumor-homing of T_EMRA_ neopeptide-specific T cells, highly represented in NCB patients (Figure 6B). Higher CXCR3 expression in T_EM_ supported the enriched effector and cytotoxic signatures found in the tumors of CD8 responders (Figure S4B). An inverse correlation between CD45RA and CD226 (Figure 6F) further supported lower functional capacity at later differentiation stages^44^. Additionally, higher CD28 and CD226 expression among T_EM_ tetramer^+^ cells (Figure 6F), i.e., those expressing lower levels of CD45RA and prevalent in CB patients (Figure 6B), supported their observed in vivo expansion under anti–PD-(L)1 therapy^26,45^.

Together, these findings demonstrate that neopeptide-specific CD8 T cells detected ex vivo exhibit memory, cytotoxic, and activation features, along with an intermediate exhaustion phenotype. Their preferential expansion under anti–PD-(L)1 therapy in clinical responders, in line with their phenotypic profile, underscores their contribution to therapeutic efficacy.

## Discussion

In this study, we systematically assessed the functional landscape of circulating T cell responses to a large set of patient-specific predicted neopeptides in advanced NSCLC. Using in vitro stimulation and ex vivo tetramer staining, we identified circulating neoantigen-specific CD8 T cells with late memory differentiation, effector function, and the ability to recognize endogenously processed peptides. The magnitude and breadth of these responses were greater in patients who experienced clinical benefit from anti–PD-(L)1 therapy. CD4 T cell responses were detectable in a larger proportion of patients and did not differ according to clinical outcome. CD8 responder status, defined by the presence of detectable neoantigen-specific CD8 T cell responses, and accompanied by CD4 T cell responses in virtually all cases, was associated with improved PFS and OS. The presence of neoantigen-specific CD8 T cells occurred independently of TMB or predicted TNB and, together with their expansion during therapy, pointed to clinical benefit being more closely linked to the engagement of functional antitumor T cell responses than to the genomic potential for neoantigenicity alone.

T cell analyses were conducted primarily in the circulating compartment, offering a unique opportunity to comprehensively evaluate tumor-specific T cell responses in a large patient cohort. The relevance of neoantigen-specific CD8 T cells identified in our assays was reinforced by their detection ex vivo using pHLA-I tetramers, which also confirmed their memory phenotype. Specific CD8 T cells were in addition able to recognize endogenously processed neoantigens. We further showed the presence of circulating neopeptide-specific clonotypes at the tumor site and the association of the circulating CD8 T cell responses with tumor immune signatures. These findings underscore the relevance of the peripheral compartment for assessing tumor-specific T cells and align with previous studies reporting overlap between circulating and tumor-infiltrating T cell repertoires^25,46^. The peripheral repertoire has been proposed as a source of TCRs or T cells for adoptive transfer therapies^32,47^. Our results further extend its utility to the investigation of mechanisms underlying response to ICB.

Most studies assessing neoantigen-specific T cell responses under anti–PD-(L)1 have focused on tumor samples, while circulating T cells have been evaluated in only a few patients, often indirectly through clonotype tracking or tetramer staining without deep phenotyping^16,24,25^. Our results demonstrate that neoantigen-specific CD8 T cells expand significantly in vivo at early timepoints, after two doses of anti–PD-(L)1, in patients with clinical responses, despite their advanced differentiation status. Tetramer^+^ cells were predominantly T_EM_ and T_EMRA_, with little to no T_CM_. These findings challenge the prevailing model that precursor exhausted T cells, typically located in the T_CM_ or stem-like memory compartments, are the sole contributors to response to ICB^30,48^. Despite their advanced differentiation, neoantigen-specific CD8 T cells expressed PD-1 and TIGIT but generally lacked CD39, consistent with an intermediate exhaustion state^26,27,49^. This was further supported by CX3CR1 expression, described in intermediate exhausted T cells retaining proliferative capacity^41,42^, in a subset of tetramer^+^ cells. In addition, T_EM_ tetramer^+^ cells expressed CD226 and CD28, markers associated with effective TCR signaling^44^ and proliferative competence under anti–PD-(L)1 therapy, given that PD-1 engagement impairs both CD28- and CD226-mediated co-stimulation^45,50,51^. In NSCLC, CD226 expression at the tumor site has been associated with improved survival following anti–PD-(L)1 therapy across three clinical trials^51^. T_EM_ tetramer^+^ cells, which expressed higher levels of CD28 and CD226, were associated with clinical outcome as higher T_EM_/T_EMRA_ ratios were found in clinical responder compared to non-responder patients. Nonetheless, we cannot exclude the presence of less-differentiated, clonally related cells in lymphoid organs may have contributed to the expanded circulating tetramer^+^ populations observed on therapy^46,52,53^. Expression of HLA-DR and CXCR3 indicated that circulating neoantigen-specific T cells were activated and retained migratory capacity toward the tumor site^54^. Finally, tetramer^+^ cells exhibited high perforin and granzyme B content, consistent with cytotoxic potential^55^, which was corroborated by in vitro degranulation assays using clonal neoantigen-specific populations.

The development of personalized neoantigen vaccines^56–60^, now beginning to show clinical efficacy^56,57^, has been driven by the growing evidence of the association between TMB and clinical responses to ICB^13–15^. Designing vaccines that elicit coordinated CD4 and CD8 T cell responses, with CD8 T cells capable of potent antitumor activity, remains challenging. Critical obstacles include the accurate selection of mutations predicted to generate immunogenic neopeptides and the optimization of vaccine platforms and formulations capable of priming integrated CD4 and CD8 responses^61^.

In our cohort, unsupervised clustering of tumors based on immune-related gene signatures segregated CD8 responders from non-responders. Tumors from CD8 responder patients were enriched in transcripts linked to antigen processing and interferon signaling, consistent with the higher stringency required for effective CD8 T cell priming compared to CD4 T cells^62^. This observation was further supported by the enrichment in tumors from CD8 responders of gene signatures comprising IFN-γ pathway components and immunoproteasome subunits, consistent with a previous report^38^. Circulating neoantigen-specific CD4 T cells were detected in virtually all CD8 responders, who constituted the group with the greatest survival benefit. Tumors from CD8 responders were also enriched in activated memory CD4 T cell signatures. Together, these findings highlight the importance of coordinated CD4 and CD8 T cell responses for optimal tumor control^63^, align with growing evidence that CD4 T cells contribute not only to CD8 T cell priming but also to the effector phase during natural immunosurveillance and immunotherapy^31,62–64^, and support the development of vaccines designed to prime such coordinated responses.

While optimizing CD4 T cell priming through vaccination should remain a priority, the more complex challenge of eliciting effective CD8 T cell responses warrants dedicated efforts. In our study, 4.4% of mutations assessed for the immunogenicity of derived predicted CD8 neopeptides tested positive. These proportions are in line with published studies assessing spontaneous T cell responses^24,40^. Compared to non-immunogenic neopeptides, immunogenic CD8 neopeptides identified in our study were predicted to bind more strongly to HLA, form more stable pHLA-I complexes, and undergo more efficient proteasomal processing. These features are consistent with prior reports^34,61^ and validate the predictive performance of HLApollo^36^, while also underscoring the biological relevance of circulating T cell responses. In contrast to earlier studies that tested smaller peptide sets^34^, we found that variant allele expression was more strongly associated with immunogenicity than overall gene expression. This distinction suggests that variant allele expression could serve as a more informative criterion for selecting mutations for inclusion in personalized vaccine platforms^61^. Nonetheless, the major future challenge in selecting mutations for vaccine design lies in predicting the immunogenicity of the encoded neopeptides, rather than focusing solely on their presentation. Compared to non-immunogenic ones, the immunogenic neopeptides identified experimentally in our study exhibited better immunogenicity prediction scores using PRIME^65^, underscoring the utility of such software for mutation prioritization in vaccine design. Importantly, data from immunogenicity studies such as ours can inform the refinement of current prediction pipelines and drive the development of new approaches.

In summary, we showed a consistent association between clinical benefit and CD8 T cell responses, whether defined by responder status, magnitude, or breadth. Moreover, CD8 responder status, which was associated with CD4 responsivness, was predictive of both PFS and OS. These associations remained robust despite the heterogeneity of the cohort and were more strongly linked to clinical outcome than TMB or predicted TNB. Notably, this prospective study was conducted in a real-world setting, with patients receiving anti–PD-(L)1 therapy as standard of care across treatment lines, with or without chemotherapy, further underscoring the relevance of neoantigen-specific T cell responses as key mediators of therapeutic efficacy and supporting the implementation of neoantigen-based vaccine and ICB combination strategies.

## METHODS

### Patient cohort and sample collection

Twenty-seven patients with locally advanced or metastatic non-small cell lung cancer (NSCLC) treated with anti–PD-1 (*n* = 18, including 5 who also received chemotherapy with either carboplatin/pemetrexed or carboplatin/taxol) or anti–PD-L1 (*n* = 9) within standard-of-care protocols at the Institut Universitaire du Cancer de Toulouse (IUCT, Toulouse, France) were prospectively enrolled in the MINER study (NCT03514368). Diagnostic tumor biopsies were collected prior to the initiation of immunotherapy and processed as formalin-fixed paraffin-embedded (FFPE) samples. Longitudinal peripheral blood samples were collected pre-treatment (T0), after 2 (T1) and 4 (T2) doses of anti–PD-(L)1, and at disease progression (T3). Peripheral blood mononuclear cells (PBMCs) were isolated by Ficoll-Hypaque density-gradient centrifugation (Sigma-Aldrich) and cryopreserved in fetal bovine serum (FBS, Gibco) supplemented with 10% dimethyl sulfoxide (DMSO, Sigma-Aldrich). Samples were collected in accordance with the Declaration of Helsinki and following Institutional Review Board approval and written informed consent. Clinical responses were evaluated using RECIST v1.1 criteria.

### DNA and RNA extraction, whole-exome and transcriptome sequencing, and data analysis

Fifteen 4 μm sections were obtained from FFPE tumor tissue blocks. One section was stained with hematoxylin and eosin, eight were used for DNA extraction (FFPE DNA Extraction Kit, Omega Bio-Tek) on a KingFisher platform, and six were used for RNA extraction (High Pure FFPET RNA Isolation Kit, Roche). All extractions from FFPE tumor tissue were performed at CellCarta (Belgium). Germline DNA was extracted from PBMCs using the QIAamp DNA Mini Kit (QIAGEN).

Whole-exome sequencing (WES) and RNA sequencing (RNA-seq) were conducted at Q² Solutions (Durham, NC). For WES, libraries were sequenced on the Illumina platform with a 100 bp paired-end protocol, at average coverage of 50X for germline and 100X for tumor DNA. RNA-seq libraries were sequenced with 50 bp paired-end reads, generating an average of 40 million reads per sample.

WES reads were aligned to the human reference genome (GRCh38/hg38) using BWA-MEM v0.7.17^66^. PCR duplicates were marked with Picard MarkDuplicates (v2.20.7), and base quality scores were recalibrated with GATK v4.1.9.0 using BaseRecalibrator and the dbSNP database (version 20180418). Somatic variants were called using three independent algorithms: Mutect2 (GATK v4.1.9)^67^, Strelka v1.0.15^68^, and LoFreq v2.1.5^69^. For LoFreq, indel quality scores were added using lofreq indelqual, and variant calling was performed using lofreq somatic with the call-indels flag. Variants passing internal quality filters for each software were retained. In addition, mutational signature profiles were analyzed with SigProfiler^70^.

HLA class I and II alleles were inferred from WES data using HLA-HD v1.3.0^35^. In cases of ambiguity, classical HLA typing was performed on germline DNA using the Miafora NGS MFLEX 11 kit (Werfen) and sequenced on an Illumina MiSeq.

RNA-seq reads were aligned to GRCh38 using STAR v2.7.5a^71^ with default parameters. Gene-level expression was quantified using HTSeq-count v0.9.1^72^ and converted to transcripts per million (TPM). For mutation-level expression, variant (alternative) and wild-type (reference) RNA reads were quantified using rnacounts^73^, which parses mutation-overlapping reads using sequence and CIGAR string information to assign allelic identity.

Tumor RNA-seq data were also used to assess immune-related gene expression profiles. Heatmaps were generated using the ComplexHeatmap v2.20.0 R package^74^. Pathway-specific immune signatures were retrieved from the Reactome database: MHC-I antigen presentation (R-HSA-983169), MHC-II antigen presentation (R-HSA-2132295), T cell receptor (TCR) signaling (R-HSA-202403), B cell receptor (BCR) signaling (R-HSA-983705), and interferon signaling (R-HSA-913531). Gene signature scores were computed using the gsva function from the GSVA package v2.2.0^75^, applying the single-sample gene set enrichment analysis (ssGSEA) method. Signature gene sets were manually curated based on literature and the CIBERSORT database^76^ (CD8 effector: *CD8A*, *EOMES*, *GZMA*, *GZMB*, *IFNG*, *PRF1*, *TBX21*; Cytotoxicity phenotype: *GZMA*, *GZMB*, *GZMK*, *IFNG*, *PRF1*; CD4 memory activated: *CCL20*, *CD2*, *CD247*, *CD28*, *CD3D*, *CD3G*, *CD40LG*, *CD6*, *CD7*, *CDC25A*, *CSF2*, *CTLA4*, *CXCL13*, *DPP4*, *GPR171*, *GPR19*, *GZMB*, *ICOS*, *IFNG*, *IL12RB2*, *IL17A*, *IL26*, *IL2RA*, *IL3*, *IL4*, *IL9*, *LAG3*, *LCK*, *LTA*, *NKG7*, *ORC1*, *PMCH*, *RRP9*, *SH2D1A*, *SKA1*, *TNFRSF4*, *TNIP3*, *TRAC, TRAT1*, *UBASH3A*; Immunoproteasome: *IFNG*, *IRF1*, *PSMB8*, *PSMB9*, *PSMB10*, *PSME1*, *PSME2*, *STAT1*; IFN- γ^77^; type I IFN^78^).

Tumor mutation clonality was inferred from WES data using ASCAT v3.1.3^79^ followed by DPClust v2.2.8^80,81^. ASCAT was used to estimate copy-number alterations, tumor ploidy, and purity (minimum read count = 10, penalty = 70, gamma = 1). DPClust was then used to cluster point mutations (PMs) by cancer cell fraction (CCF) (minimum read depth = 10 and minimum alt depth = 2); insertions and deletions (indels) were not considered for clustering. If the cluster with the highest CCF did not fall between 0.95 and 1.05, ASCAT was rerun using adjusted values for purity (rho_manual) and ploidy (psi_manual). The adjusted purity was computed as: purity × CCF of the dominant cluster. Ploidy was re-estimated using the reestimate_ploidy function (https://github.com/tlesluyes/myFun). DPClust was then rerun with the updated ASCAT profile. For each tumor, the clonal cluster was defined as that with a CCF of 1. Indels were classified as clonal or subclonal based on their variant allele fraction relative to the sample’s estimated subclonal threshold. Tumors with a purity < 20% were not considered for clonality analyses.

### Neopeptide prediction, selection and synthesis

Somatic mutations identified by at least two variant-calling algorithms were annotated using VEP v106^82^. Exonic, non-synonymous mutations were retained, excluding those causing loss of the start codon or gain of a stop codon. Selected variants included missense PMs, in-frame indels (IFs), and frameshift indels (FSs). Expression filtering was applied based on RNA-seq data. For mutated positions covered by ≥ 20 RNA reads, mutations were retained if ≥ 2 variant reads were detected. For mutated positions with < 20 reads, gene-level expression was used as a proxy, with a threshold ≥ 30 TPM^34^.

Sequences harboring expressed missense PMs and IFs were translated into at least 29-amino acid sequences centered on the PM or indel. Sequences containing FSs and stop lost PMs were translated starting from 14 codons upstream of the variant site and until a stop codon was encountered. The resulting sequences were used to predict neopeptides capable of binding to each patient’s HLA-I and HLA-II alleles. Presentation prediction for 9–11-mer neopeptides by patient-specific HLA-I alleles were performed using HLApollo^36^. For each patient, the top 210 or 420 predicted HLA-I binders (ranked by HLApollo score) were synthesized. Two peptide pools of 210 neopeptides each were generated: Pool 1 comprised the top 210 binders; Pool 2 contained the next 210. Each pool was deconvoluted into 21 minipools (MPs) of 20 peptides, using a matrix design with one peptide shared between two MPs to allow response deconvolution^39^.

For HLA-II, the binding of 9–15-mer neopeptides to patient-specific alleles was predicted using netMHCIIpan v4.0^83^. Peptides with binding scores ≤ 10% were retained, as previously recommended^83^. For neopeptides derived from PMs or IFs, all predicted binders sharing the same mutation were aligned to generate an extended peptide sequence (Figure S1A and S1B). If the extended sequence was ≤ 24 amino acids, it was synthesized as is (Figure S1A). Sequences of 25–26 amino acids were divided into two overlapping 20-mers and sequences > 26 amino acids were covered by 20-mers overlapping by ≥ 15 amino acids (Figure S1B). For FS mutations, 20-mers overlapping by at least 10 amino acids were synthesized to span the translated product (Figure S1C). For each patient, neopeptides covering up to 50 mutations with the best HLA-II binding predictions were selected. The corresponding long peptides were synthesized and pooled into a single pool per patient, which was deconvoluted into 11 MPs using the same matrix approach as above^39^.

All single peptides, pools, and MPs were produced by GenScript and dissolved in DMSO. Once the immunogenicity of predicted neopeptides was assessed, immunogenic and non-immunogenic HLA-I–restricted neopeptides were compared using additional algorithms. Binding affinities were predicted using netMHCpan v4.0^83^ and MixMHCpred v3.0^84^. The stability of peptide–HLA-I complexes was assessed using netMHCstabpan v1.0^85^, and the likelihood of proteasomal cleavage was predicted using netChop v3.1 for both standard proteasome (20S) and immunoproteasome (C-term)^86^. The probability of TCR recognition was evaluated with PRIME v2.1^65^.

### In vitro stimulation, assessment of T cell responses, and isolation and cloning of specific T cells

CD8, CD14, and CD4 cells were sequentially isolated by magnetic cell sorting (Miltenyi Biotec) from mixed pre-treatment and on-treatment PBMCs of each patient. CD8 or CD4 T cells were stimulated with neopeptide pools (1 μM/peptide) in the presence of autologous CD14 monocytes as antigen-presenting cells (APCs) at a 4:1 T cell:monocyte ratio. Cultures were maintained in Iscove’s Modified Dulbecco’s Medium (IMDM, Sigma-Aldrich) supplemented with 1% MEM non-essential amino acids (Gibco, Thermo Fisher Scientific), 1% penicillin–streptomycin (Sigma-Aldrich), 4% L-glutamine (Gibco), 10% human serum (Institut de Biotechnologies Jacques Boy), and recombinant human (rh) IL-2 (50 IU/mL), rhIL-7 (10 ng/mL), and rhIL-15 (2.5 ng/mL) (all Miltenyi Biotec). CD8 T cells were also stimulated with a control pool of 32 viral epitopes derived from CMV, EBV, and influenza (CEF pool; 0.6 μM/peptide, Miltenyi Biotec).

On day 11, cultures were stimulated or not with the same peptide pools for 4 hours. Brefeldin A (3 μg/mL, eBioscience, Thermo Fisher Scientific) was added 1 hour after the start of stimulation. Cells (2×10⁵) were stained for viability (Fixable Viability Dye eFluor™ 506, eBioscience) in PBS for 20 minutes at 4°C. Surface staining was performed using fluorochrome-conjugated monoclonal antibodies (mAbs) against CD3 (UCHT1, BUV395, BD Biosciences) and CD8 (RPA-T8, BV421, BD Biosciences) or CD4 (RPA-T4, BV421, BD Biosciences) in PBS with 5% FBS (staining buffer) for 15 minutes at 4°C. Cells were then washed, fixed (2% glucose, 1% paraformaldehyde in PBS) for 10 minutes at room temperature, permeabilized with 0.1% saponin in staining buffer, and stained intracellularly with mAbs against IFN-γ (B27, AF700, BD Biosciences) and TNF-α (MAb11, PE-Cy7, BD Biosciences) for 30 minutes at room temperature. Flow cytometry was performed on a BD LSRFortessa X20, and data were analyzed using BD FACSDiva (BD Biosciences) and FlowLogic™ (Miltenyi Biotec).

Positive cultures were further assessed by intracellular cytokine staining after stimulation with neopeptide MPs (day 12) and single peptides (day 14). Negative cultures were re-stimulated with the same pool of neopeptides in the presence of CD14 monocytes, and day 7–10 cultures were assessed for reactivity to pools, MPs, and individual peptides.

In parallel, neopeptide responder patients were tested using neopeptide pools composed exclusively of immunogenic neopeptides. These cultures were evaluated for recognition of mutated and wild-type peptide counterparts by intracellular cytokine staining as above. Cultures were also used for the isolation of specific T cells via IFN-γ secretion assay (Miltenyi Biotec). Briefly, cultures were stimulated or not with the immunogenic neopeptide pool (1 μM/peptide) for 1 h45, washed in PBS with 0.5% BSA and 2 mM EDTA (staining buffer 2), and incubated with IFN-γ catch reagent on ice. Cells were transferred to 10-40 mL of culture medium at 37°C for 45 minutes under continuous rotation, washed, and stained with PE-conjugated IFN-γ detection antibody and surface mAbs against CD3, CD8 or CD4 (as above) for 10 minutes on ice. IFN-γ–secreting cells were sorted using a FACSAria Fusion (BD Biosciences). Sorted cells were either snap-frozen in liquid nitrogen as dry pellets for TCR sequencing or cloned by limiting dilution in Terasaki plates (Greiner Bio-one) in the presence of 35 Gy-irradiated allogeneic PBMCs (Etablissement Français du Sang, Toulouse), PHA-L (1 μg/mL, Sigma-Aldrich), and rhIL-2 (150 IU/mL). Clones were expanded by periodic stimulation under identical conditions.

### Functional characterization of CD8 clones

The specificity of CD8 T cell clones was first assessed by testing their ability to recognize individual peptides from the pool of immunogenic neopeptides used for their isolation. Clones were cultured overnight with individual neopeptides (1 μM/peptide) in the presence of rhIL-2 (20 IU/mL), and IFN-γ secretion in the supernatant was quantified by ELISA (Invitrogen, Thermo Fisher Scientific).

Recognition of the wild-type (WT) peptide was then evaluated. Autologous CD14 monocytes (1.5 × 10⁴) were pulsed for 3 hours with either neopeptides or their WT counterparts (10 μM/peptide), washed, and co-cultured overnight with CD8 clones (5 × 10³ cells). IFN-γ and granzyme B (GzmB; R&D Systems) were measured in the supernatant by ELISA.

To test the ability of clones to recognize endogenously processed neoantigens, tandem minigenes (TMGs) were designed as described^87^. For each patient, TMGs were designed to encode 27-mer peptides encompassing all patient-specific mutations identified as encoding immunogenic neopeptides, with the mutant amino acid positioned at residue 14. TMGs were codon-optimized and cloned into pcDNA3.1(+) using BamHI and EcoRI sites (GenScript). COS-7 cells (1.5 × 10⁶) were co-transfected, with the TMG and a plasmid encoding the relevant HLA-I allele under the EF1a promoter^36,88^, using electroporation (Amaxa 4D-Nucleofector, CM-130 program, Lonza). COS-7 cells were cultured in Dulbecco’s Modified Eagle Medium (DMEM, Sigma-Aldrich) supplemented with 10% FBS and 1% penicillin–streptomycin. Antibiotics were removed from the culture medium during the 24 hours preceding electroporation.

After 24 hours, COS-7 cells were detached using Trypsin-EDTA (Sigma-Aldrich) and used as target cells in co-culture with CD8 clones at a 2:1 effector-to-target ratio. Co-cultures were performed in IMDM with 10% human serum and rhIL-2 (20 IU/mL), in the presence or absence of the corresponding immunogenic neopeptide (10 μM/peptide) and fluorochrome-conjugated anti-CD107a mAb (H4A3, APC, BD Biosciences). After 4 hours, cells were stained with viability dye and mAbs against CD3 and CD8 (as described above), and CD107a surface expression was analyzed by flow cytometry. At 24 hours, additional staining for 4-1BB (4B4-1, PE, BD Biosciences) was performed, and IFN-γ and GzmB levels were measured by ELISA in the supernatants.

### CDR3 sequencing of neopeptide-specific circulating T cells and alignment with tumor RNA-seq data

DNA was extracted from IFN-γ–producing, neopeptide-specific circulating CD8 and CD4 T cells isolated from in vitro-stimulated cultures of 12 patients, as detailed above. Deep sequencing of the complementarity-determining region 3 (CDR3) of the TCRβ chain was performed (Adaptive Biotechnologies). The number of sequenced neoantigen-reactive T cells per sample ranged from 4.3 × 10³ to 6.7 × 10⁴. Productive TCRβ rearrangements were used to quantify clonotype frequencies for CD8 and CD4 T cells.

Circulating CDR3 sequences from all patients were compiled into a FASTA-format reference (CDR3-reference). Each CDR3-reference was aligned using blastn v2.16.0 (default parameters) to the V and J exons of the human genome, in order to annotate the final nucleotide of the V region and the initial nucleotide of the J region for each CDR3. Tumor RNA-seq reads either mapped to chromosome 7 or unmapped to any chromosome were aligned against the CDR3-reference using blastn with default settings. Alignments were retained according to the following criteria: (i) at least 5 nucleotides upstream and downstream of the annotated V–N1 and/or N2–J junctions were aligned and (ii) no gaps or mismatches were allowed within the aligned read/CDR3 sequences. Among 125 candidate RNA reads, alignments were manually validated if they spanned at least one full VD or DJ junction without mismatches to the CDR3 sequence.

### pHLA-I tetramer production and phenotypic assessment of neoantigen-specific CD8 T cells

For each patient, all possible combinations between immunogenic 9- to 11-mer neopeptides and that patient’s HLA-I alleles were evaluated, and those with an HLApollo score ≤ 0.05 were considered to generate pHLA-I tetramers. Biotinylated HLA-I monomers harboring a UV-cleavable peptide (UV-HLA-I) corresponding to 31 alleles were available for tetramer production. Peptide exchange reactions were performed, as previously described^89^, in 96-well UV-Star plates (Greiner Bio-One) by incubating UV-HLA-I (1.08 µM final concentration) with the relevant neopeptide (100-fold molar excess) and 5% (v/v) ethylene glycol (Sigma-Aldrich) in exchange buffer (25 mM Tris pH 8.0, 150 mM NaCl, 2 mM EDTA). Plates were exposed to UV light (365 nm) for 25 minutes at room temperature and then sealed and incubated overnight. Exchange mixtures were incubated with PE-labeled streptavidin (BioLegend) at a pHLA-I:streptavidin ratio of 6:1 in v-bottom 96-well polypropylene plates (Corning). Tetramers were centrifuged at 1,850 rpm for 5 minutes to remove aggregates. Validation of tetramer functionality was performed by staining CD8 T cell cultures stimulated with the immunogenic neopeptide pool. Cells (5 × 10⁴) were stained with a viability dye (Fixable Viability Dye eFluor™ 506, Thermo Fisher Scientific) for 10 minutes at room temperature, washed, and incubated with tetramer for 30 minutes at 4°C, followed by staining with anti-CD3 (UCHT1, BUV737, BD Biosciences) and anti-CD8 (RPA-T8, BUV496, BD Biosciences) mAbs. Cells were then fixed and analyzed by flow cytometry. Specific CD8 T cell clones were stained under the same conditions.

For ex vivo phenotyping, CD8 T cells were positively selected from pre-treatment (T0) and on-treatment (T1) PBMCs using magnetic sorting. Tetramer staining was performed as above, followed by surface staining with fluorochrome-conjugated mAbs specific for: CD3 (UCHT1, BV650 or BV421), CD8 (RPA-T8, BUV496), CD45RA (HI100, Invitrogen), CCR7 (150503, PerCP-Cy5.5, BD Biosciences), CD28 (CD28.2, BUV395), PD-1 (EH12.1, BUV737), TIGIT (MBSA43, PECy7, Invitrogen), CXCR3 (1C6/CXCR3, APC), HLA-DR (LN3, APC-eFluor780), CD57 (QA17A04, BV605, BioLegend), CD226 (11A8, BV785), CX3CR1 (2A9-1, APC-Cy7), CXCR5 (RF8B2, RB780), and CD39 (TU66, BV650). Intracellular staining was performed for granzyme B (GB11, AF700) and perforin (δG9, AF647). Cells were fixed and analyzed using a BD LSRFortessa X20 cytometer. Data were analyzed using BD FACSDiva, FlowJo v10.10.0 and Flowlogic v7.3. Uniform Manifold Approximation and Projection (UMAP) analyses were conducted on tetramer⁺ cells based on the fluorescence intensity of all phenotypic markers, excluding viability dye, tetramers, and PD-1.

### Statistical analyses

For correlation and survival analyses involving tumor mutational burden (TMB) or tumor neoantigenic burden (TNB), normalization was performed across all 27 tumor WES datasets by including only mutations located in regions with coverage ≥ 50 reads. Normality was assessed using the Shapiro-Wilk test. For normally distributed values, paired or unpaired t-tests were applied. Line and error bars represent the mean ± SEM. For non-normally distributed data, paired and unpaired comparisons were made using the Wilcoxon and the Mann-Whitney tests, respectively, with results shown as median ± IQR. When applicable, *P* values were adjusted with Benjamini-Hochberg method. Pearson’s correlation was used for association analyses. Categorical associations between two variables were assessed using Chi-squared or Fisher’s exact test. For survival analyses, Kaplan–Meier curves were plotted and compared using the log-rank test. Statistical results are annotated in figures as follows: ns, not significant (*P* ≥ 0.05); \**P* < 0.05; \*\**P* < 0.01; \*\*\**P* < 0.001; \*\*\*\**P* < 0.0001. Analyses were conducted using GraphPad Prism (v10), R (v4.4.3) and STATA (v18) software.

## Data Availability

WES and RNA-seq data generated in this study will be deposited in the European Genome-Phenome Archive (EGA) and accession numbers will be made available upon manuscript acceptance.

## Acknowledgments

The authors express their deep gratitude to the patients who participated in this study. We thank the *Direction de la Recherche Clinique* (*Oncopole Claudius Regaud, Institut Universitaire du Cancer de Toulouse*), sponsor and investigator of the MINER protocol, and in particular Dr. Muriel Mounier and Mrs. Muriel Poublanc for their support throughout the study. We are also grateful to Mrs. Aurore Komperdra (*Centre Hospitalier Universitaire, Institut Universitaire du Cancer de Toulouse*) for her valuable assistance with patient accrual and clinical data collection, and to Dr. Philippe Rochaix and Mrs. Laurence Puydenus (*Support biopathologique des essais cliniques, Oncopole Claudius Regaud, Institut Universitaire du Cancer de Toulouse*) for their help with tumor sample retrieval. We are grateful to the Genotoul bioinformatics platform Toulouse Occitanie (bioinfo.genotoul.fr/) for computing and storage resources. Figures 1C and 2A were created in BioRender.

This research was made possible through a collaboration within the imCORE Network, supported by F. Hoffmann-La Roche Ltd. Célia Ramade was supported through PhD fellowships from the University of Toulouse and the *Fondation ARC pour la recherche sur le cancer* as well as through the CARe Graduate School (ANR-18-EURE-0003 in the framework of the *Programme des Investissements d’Avenir*).

## Author contributions

Conceptualization: C. Ramade, L.D. and M.A.; Formal analysis: C. Ramade, N.T., C.M.S., B.C., T.F. and M.A.; Clinical investigation: J.P.D., C.G.R., A.M., J.M.; Laboratory investigation: C. Ramade, N.T., C.M.S., D.O., F.L.V., S.J., M.H., C. Fournier, A.S., M. Michelas, V.S., G.C.L, V.F., M. Maixent, L.S., M.X.H., C. Ross, H.X., T.L., N.C.J. and C.D.; Resources: M.D., A.H., J.P.D. and J.M.; Data curation: N.T., B.C. and T.F.; Writing – original draft: C. Ramade and M.A.; Writing – review & editing: C. Ramade, N.T., C.M.S., D.O., B.C., A.M., C.D., T.F., J.P.D., J.M., L.D. and M.A.; Visualization: C. Ramade, N.T., C.M.S.; Supervision: M.A.; Project administration: F.B., C. Fonseca and M.A.; Funding acquisition: M.A.

## Competing interests

D.O., S.J., M.H., M.X.H., M.D., A.H., C. Ross, H.X. and L.D. were employees of Genentech and F.B. and C. Fonseca were employees of F. Hoffmann-La Roche Ltd. at the time of the study. All other authors declare no competing interests.

---

**Table S1.**
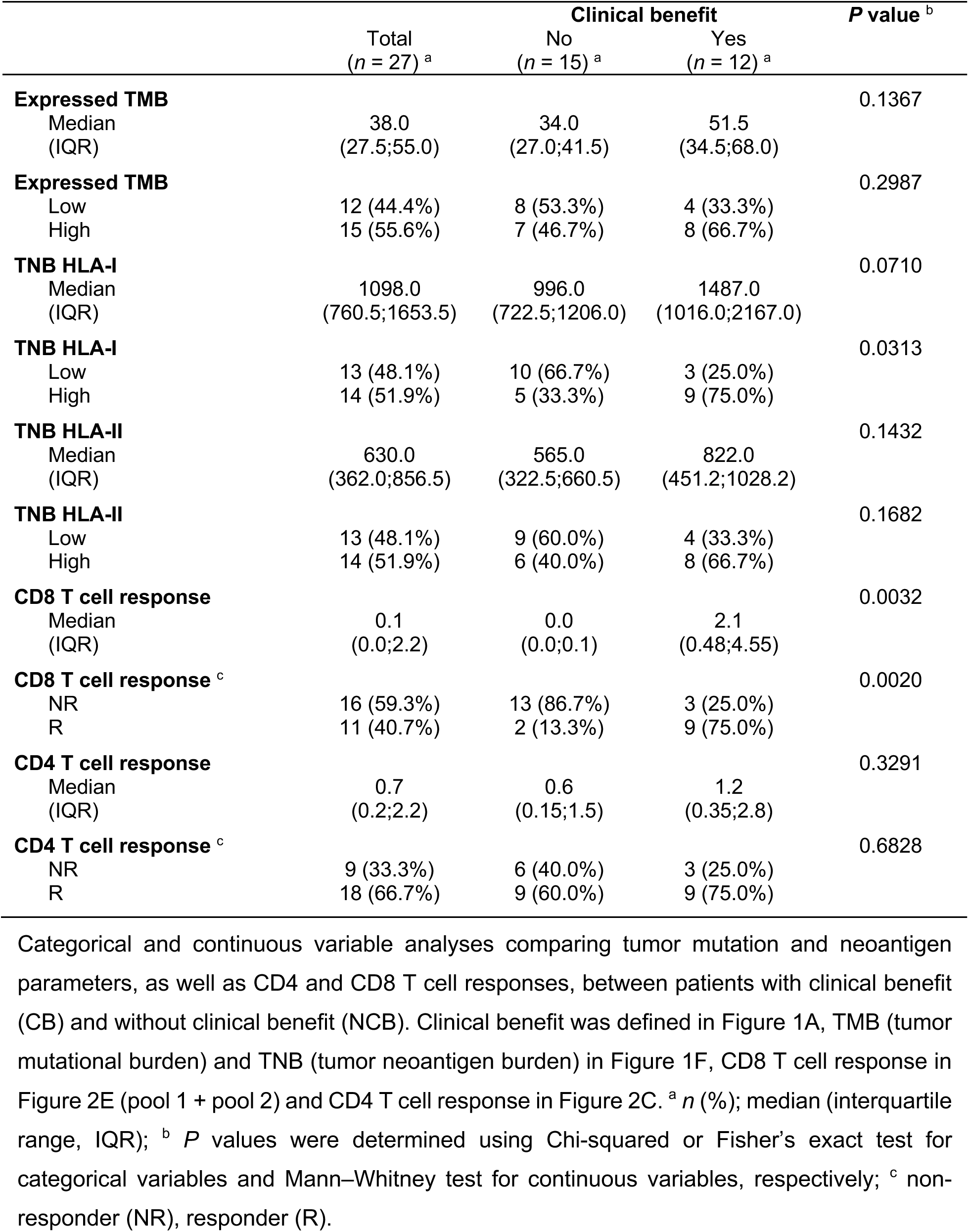
Correlation between clinical benefit and tumor mutation burden, neoantigen landscape, and T cell responses.

**Figure S1.**
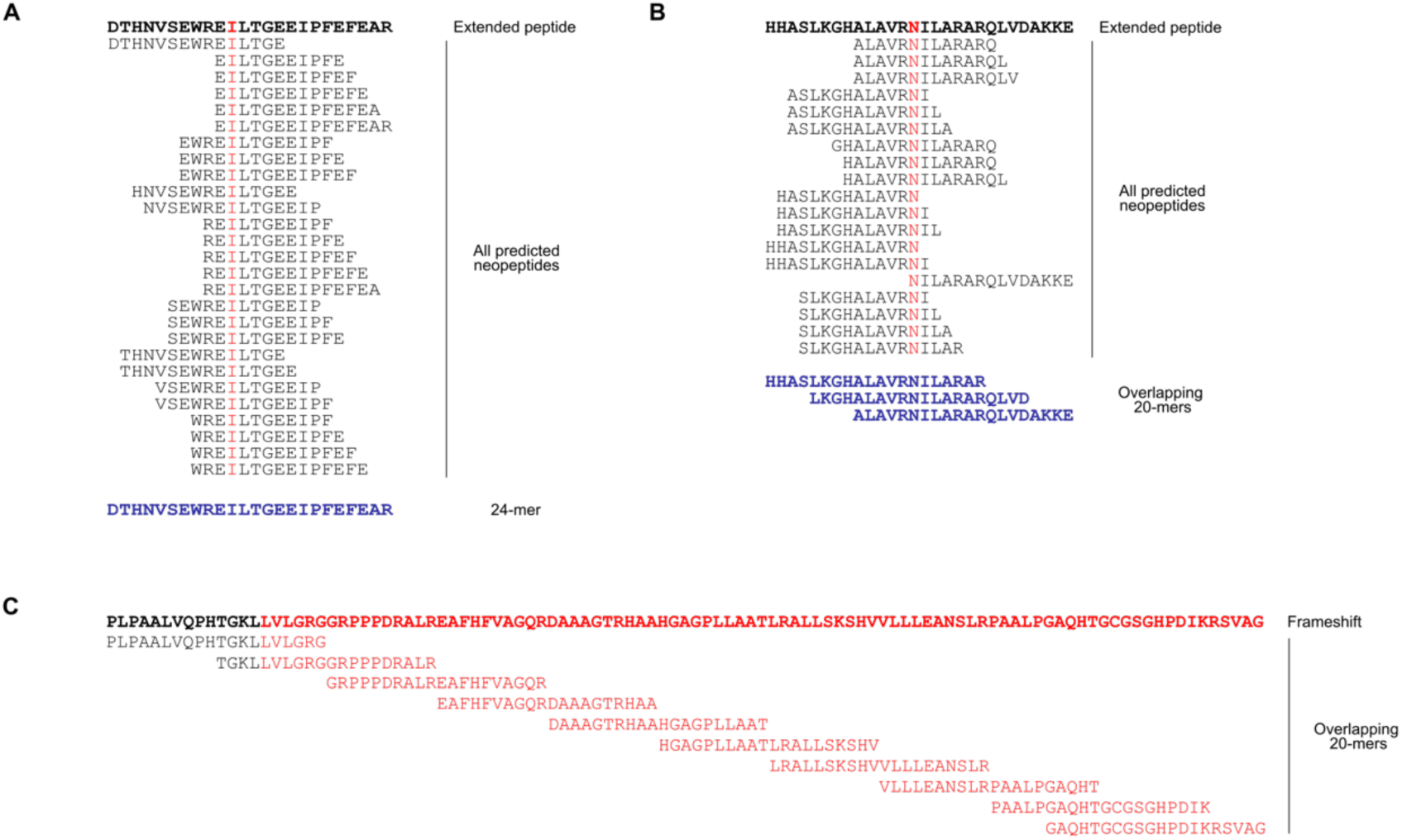
Design of long peptides used for CD4 T cell stimulation. (A,B) For each tumor variant (excluding indels leading to frameshifts), all predicted HLA-II–binding neopeptides containing the same mutation were aligned to generate an extended peptide sequence. If the resulting sequence was ≤ 24 amino acids, it was synthesized as a single peptide (blue, example in A). Sequences of 25–26 amino acids were split into two overlapping 20-mers. For longer sequences, 20-mer peptides overlapping by at least 15 amino acids were synthesized (blue, example in B). (C) Sequences containing frameshift indels were translated starting from 14 codons upstream of the variant site and until a stop codon was encountered. 20-mers overlapping by at least 10 amino acids were synthesized to span the translated product. Each patient’s CD4 peptide pool (Figure 2A) included long peptides covering 50 mutations encoding the best predicted HLA-II binders.

**Figure S2.**
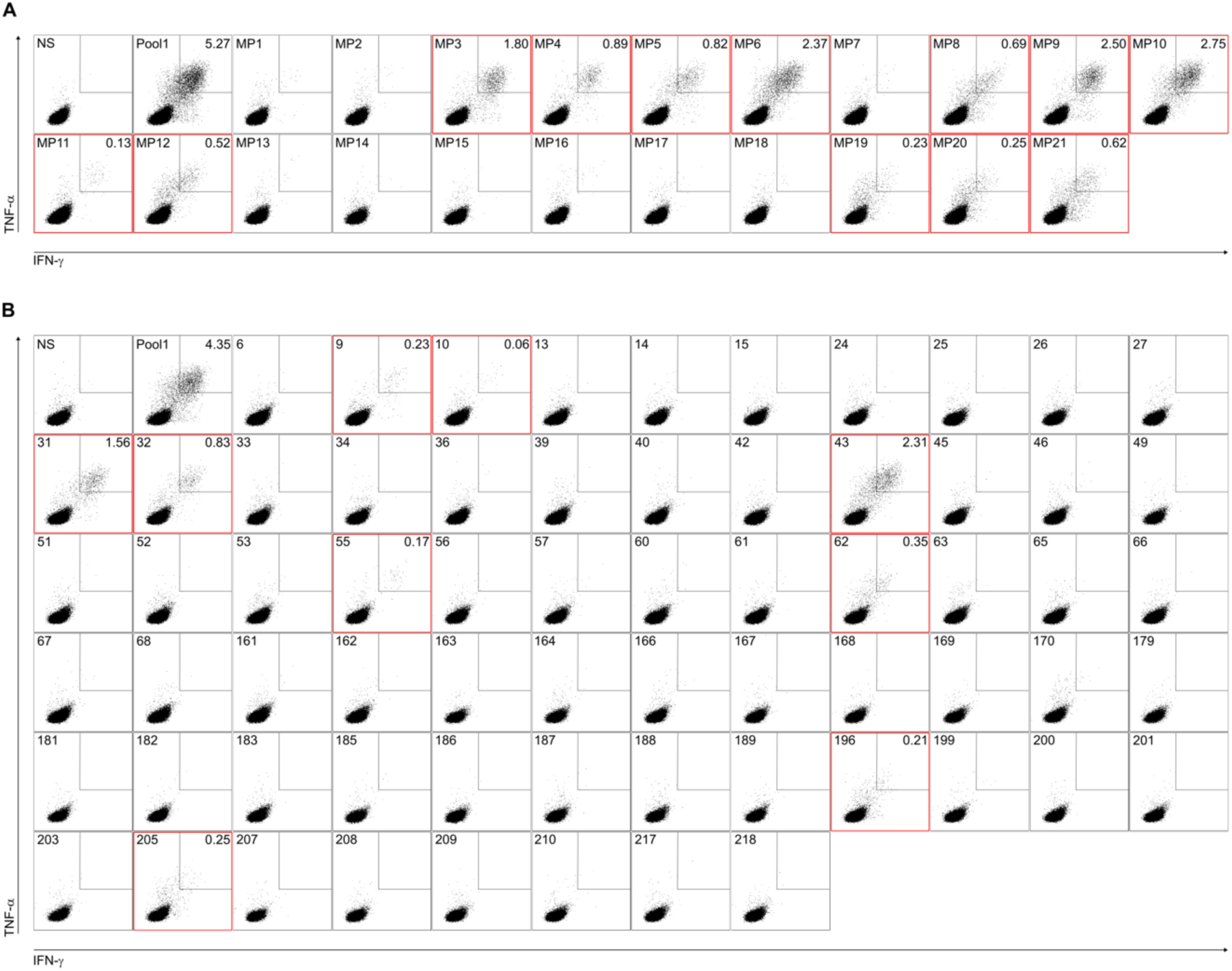
Deconvolution strategy for identification of immunogenic neopeptides recognized by CD8 T cells. Day 11 CD8 T cell cultures from in vitro stimulated (IVS)1 (Figure 2A, 2D and 2E) and IVS2 (Figure 2A) that tested positive for neopeptide pools were screened under unstimulated conditions (NS) and after stimulation the corresponding neopeptide pool or with 21 minipools (MPs) composed of 20 neopeptides each (A). Based on MPs found positive in A, shared peptides were identified and tested individually (B). Representative dot plots show IFN-γ and TNF-α production for one culture. Stimulation condition is indicated in the top left corner. Red squares denote positive responses; percentages of IFN-γ⁺TNF-α⁺ cells are shown in the top right corner of positive conditions.

**Figure S3.**
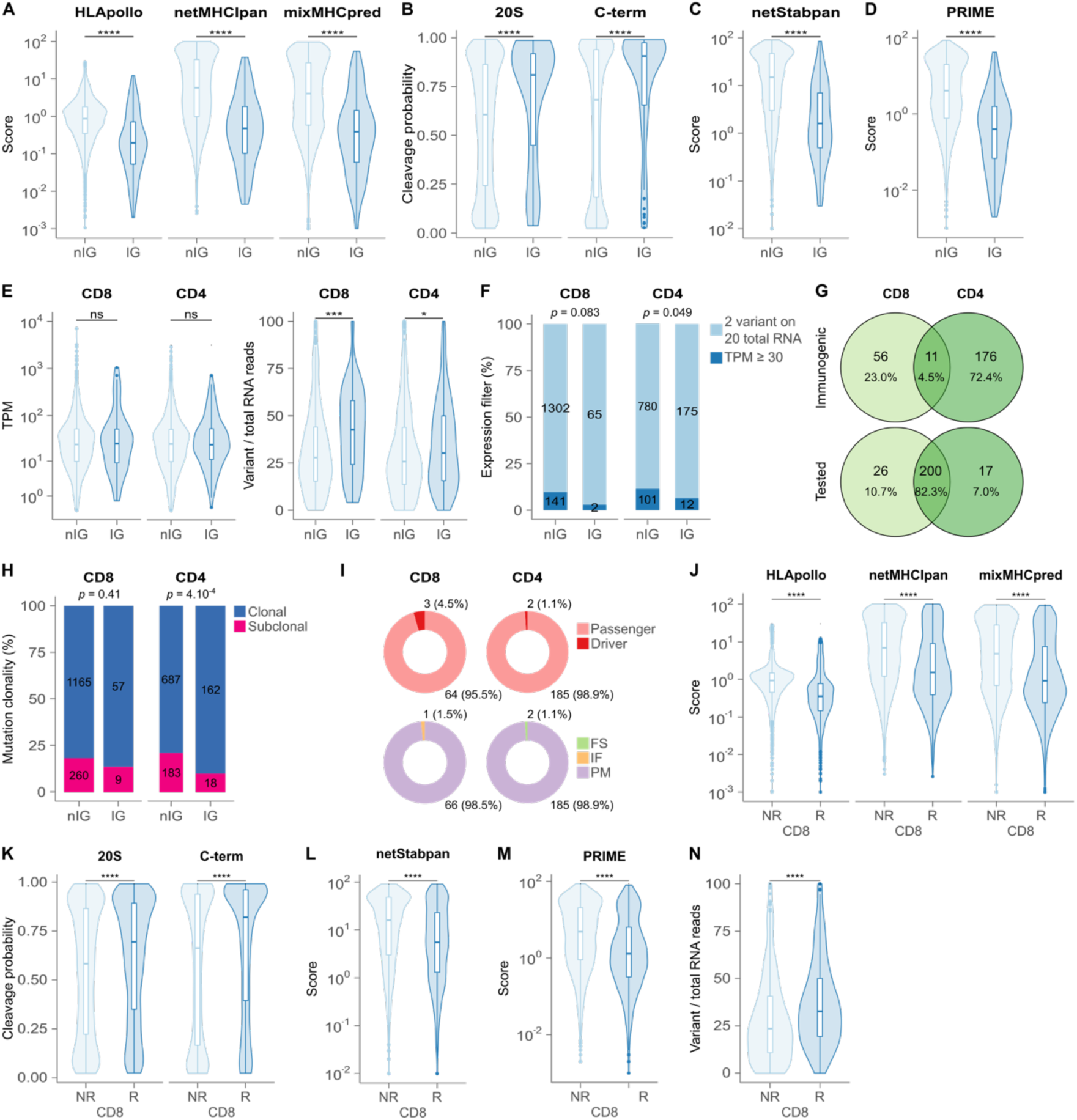
Intrinsic features of predicted and immunogenic neopeptides and of their encoding mutations. (A–I) Among 7,038 predicted CD8 neopeptides, 108 derived from 67 mutations were classified as immunogenic (IG; Figure 2H), and 6,930 neopeptides as non-immunogenic (nIG). For CD4 T cell analysis, 2,103 long peptides (Figure S1), encompassing 21,453 predicted HLA-II binders, were tested and originated from 1,068 mutations. IG CD4 neopeptides were derived from 187 mutations. (A) Binding score (HLApollo) of the best neopeptide–HLA-I pair for IG and nIG CD8 neopeptides; the same pairs were evaluated with netMHCIpan and mixMHCpred. (B) Predicted cleavage probability by the standard proteasome (20S) and immunoproteasome (C-term) (netChop). (C) Stability score of pHLA-I complexes (netStabpan). (D) Predicted TCR recognition probability (PRIME). (E) Gene expression levels (TPM; left) and ratio of variant to total RNA reads (right) for mutations encoding IG and nIG neopeptides. (F) Number and percentage of mutations encoding IG and nIG CD8 and CD4 neopeptides according to expression thresholds: ≥2 variant reads out of ≥20 total (light blue); if <20 reads, then based on gene-level TPM (Figure 1C). (G) Numbers and proportions of mutations encoding IG CD8 and/or CD4 neopeptides (top) and, for the same set of mutations, whether they were tested for CD8 and/or CD4 reactivity (bottom). (H) Distribution of clonal and subclonal mutations encoding IG and nIG neopeptides. Neopeptides of patient MIN-130 were excluded as tumor purity precluded mutation clonality analyses. (I) Among mutations encoding IG neopeptides, the proportion of driver vs passenger mutations (top), and mutation types: point mutations (PM), in-frame indels (IF), and frameshift indels (FS) (bottom). (J–N) Analysis restricted to the top 210 predicted HLA-I neopeptides per patient (pool 1; Figure 2A). This included 5,607 neopeptides derived from 1,426 mutations (2,301 neopeptides from 769 mutations in CD8 responders and 3,306 neopeptides from 657 mutations in CD8 non-responders). For each neopepitde, the best-ranked neopeptide–HLA-I pair was used for comparison between groups for binding score (J; as in A), cleavage probability (K; as in B), stability score (L; as in C), predicted TCR recognition (M; as in D), and mutant-to-total RNA ratio (N; as in E). Mann–Whitney test (A–E and J–N) and Fisher’s exact test (F and H). \**P* < 0.05, \*\*\**P* < 0.001, \*\*\*\**P* < 0.0001; *ns*, not significant.

**Figure S4.**
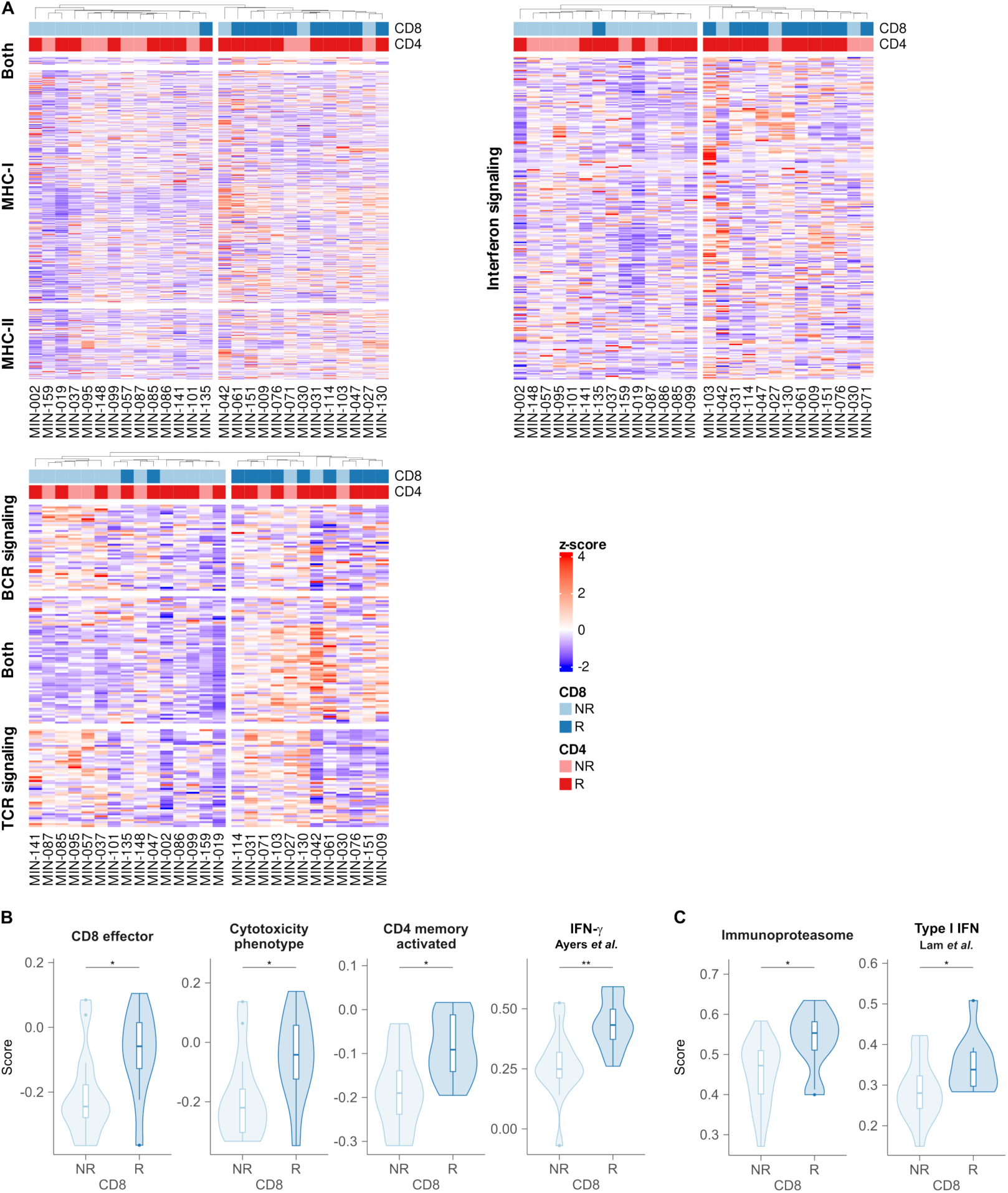
Tumor immune gene expression signatures associate with circulating neoantigen-specific CD8 T cell responses. (A) Unsupervised clustering of patients based on tumor RNA-seq gene expression (Figure 1C). Heatmaps display scaled normalized expression (z-scores) of gene sets grouped into biological pathways: antigen presentation (top left), interferon signaling (top right), and BCR/TCR signaling (bottom left). Patient annotations include CD4 and CD8 T cell responder (R) or non-responder (NR) status, as defined in Figure 2C and 2E, respectively. (B,C) Gene signature scores derived from tumor RNA-seq data for CD8 T cell responders (CD8 R, *n* = 11) and non-responders (CD8 NR, *n* = 16). Mann–Whitney test with Benjamini-Hochberg adjusted *P* values (B and C). \**P* < 0.05; \*\**P* < 0.01.

**Figure S5.**
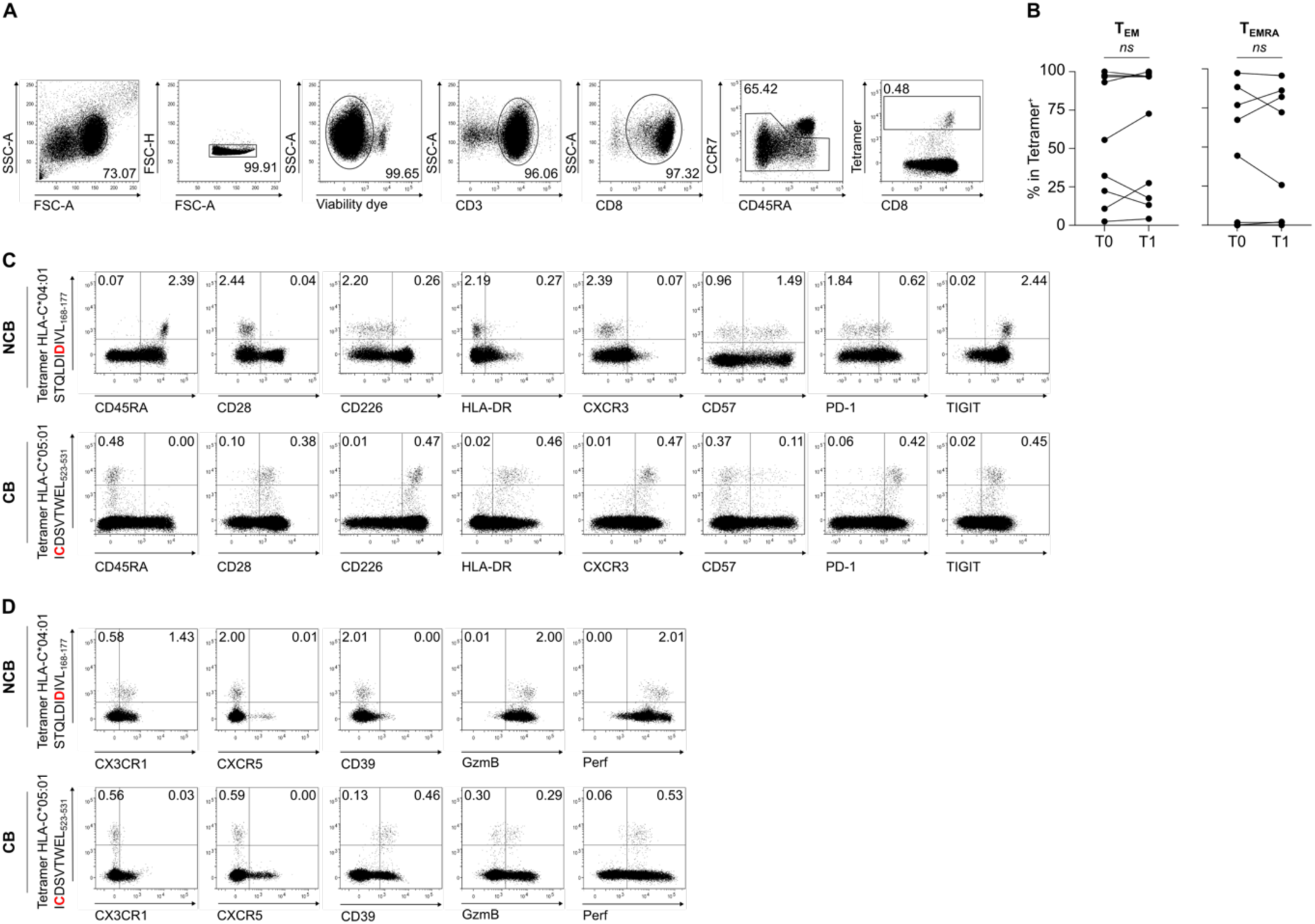
Ex vivo tetramer staining of circulating CD8 T cells. CD8 T cells isolated from baseline (T0) and on-treatment (T1) PBMCs were stained ex vivo with personalized pHLA-I tetramers and analyzed by flow cytometry. (A) Gating strategy for identifying tetramer⁺ cells. (B) Summary of memory tetramer⁺ cells distributed across effector memory (T_EM_: CCR7⁻CD45RA⁻) and T_EMRA_ (CCR7⁻CD45RA⁺) subsets, before (T0) and on therapy (T1). (C,D) Representative dot plots from patients without clinical benefit (NCB) and with clinical benefit (CB), showing tetramer staining versus the indicated phenotypic markers. Distinct monoclonal antibody panels were used in C and D, as noted. Gating was performed as shown in A. The neopeptide sequence, with the mutated position in red, and the restricting HLA-I allele are indicated. Numbers denote the percentage of cells in each quadrant. Perf, perforin; GzmB, granzyme B. Wilcoxon matched-pairs rank test (B). *ns*, not significant.

## References

1. Ready, N., Hellmann, M.D., Awad, M.M., Otterson, G.A., Gutierrez, M., Gainor, J.F., Borghaei, H., Jolivet, J., Horn, L., Mates, M., et al. (2019). First-Line Nivolumab Plus Ipilimumab in Advanced Non-Small-Cell Lung Cancer (CheckMate 568): Outcomes by Programmed Death Ligand 1 and Tumor Mutational Burden as Biomarkers. J. Clin. Oncol. 37, 992–1000. 10.1200/JCO.18.01042.

2. Reck, M., Rodríguez-Abreu, D., Robinson, A.G., Hui, R., Csőszi, T., Fülöp, A., Gottfried, M., Peled, N., Tafreshi, A., Cuffe, S., et al. (2016). Pembrolizumab versus Chemotherapy for PD-L1–Positive Non–Small-Cell Lung Cancer. N Engl J Med 375, 1823–1833. 10.1056/nejmoa1606774.

3. West, H., McCleod, M., Hussein, M., Morabito, A., Rittmeyer, A., Conter, H.J., Kopp, H.-G., Daniel, D., McCune, S., Mekhail, T., et al. (2019). Atezolizumab in combination with carboplatin plus nab-paclitaxel chemotherapy compared with chemotherapy alone as first-line treatment for metastatic non-squamous non-small-cell lung cancer (IMpower130): a multicentre, randomised, open-label, phase 3 trial. Lancet Oncol 20, 924–937. 10.1016/S1470-2045(19)30167-6.

4. O’Brien, M., Paz-Ares, L., Marreaud, S., Dafni, U., Oselin, K., Havel, L., Esteban, E., Isla, D., Martinez-Marti, A., Faehling, M., et al. (2022). Pembrolizumab versus placebo as adjuvant therapy for completely resected stage IB-IIIA non-small-cell lung cancer (PEARLS/KEYNOTE-091): an interim analysis of a randomised, triple-blind, phase 3 trial. Lancet Oncol 23, 1274–1286. 10.1016/S1470-2045(22)00518-6.

5. Forde, P.M., Spicer, J.D., Provencio, M., Mitsudomi, T., Awad, M.M., Wang, C., Lu, S., Felip, E., Swanson, S.J., Brahmer, J.R., et al. (2025). Overall Survival with Neoadjuvant Nivolumab plus Chemotherapy in Lung Cancer. N Engl J Med. 10.1056/NEJMoa2502931.

6. Tumeh, P.C., Harview, C.L., Yearley, J.H., Shintaku, I.P., Taylor, E.J.M., Robert, L., Chmielowski, B., Spasic, M., Henry, G., Ciobanu, V., et al. (2014). PD-1 blockade induces responses by inhibiting adaptive immune resistance. Nature 515, 568–571. 10.1038/nature13954.

7. Wang, S.L., and Chan, T.A. (2025). Navigating established and emerging biomarkers for immune checkpoint inhibitor therapy. Cancer Cell 43, 641–664. 10.1016/j.ccell.2025.03.006.

8. Hanada, K., Zhao, C., Gil-Hoyos, R., Gartner, J.J., Chow-Parmer, C., Lowery, F.J., Krishna, S., Prickett, T.D., Kivitz, S., Parkhurst, M.R., et al. (2022). A phenotypic signature that identifies neoantigen-reactive T cells in fresh human lung cancers. Cancer Cell 40, 479–493.e6. 10.1016/j.ccell.2022.03.012.

9. Linnemann, C., van Buuren, M.M., Bies, L., Verdegaal, E.M.E., Schotte, R., Calis, J.J.A., Behjati, S., Velds, A., Hilkmann, H., Atmioui, D. el, et al. (2015). High-throughput epitope discovery reveals frequent recognition of neo-antigens by CD4+ T cells in human melanoma. Nat Med 21, 81–85. 10.1038/nm.3773.

10. Bobisse, S., Genolet, R., Roberti, A., Tanyi, J.L., Racle, J., Stevenson, B.J., Iseli, C., Michel, A., Le Bitoux, M.-A., Guillaume, P., et al. (2018). Sensitive and frequent identification of high avidity neo-epitope specific CD8 + T cells in immunotherapy-naive ovarian cancer. Nat Commun 9, 1092. 10.1038/s41467-018-03301-0.

11. Stevanović, S., Pasetto, A., Helman, S.R., Gartner, J.J., Prickett, T.D., Howie, B., Robins, H.S., Robbins, P.F., Klebanoff, C.A., Rosenberg, S.A., et al. (2017). Landscape of immunogenic tumor antigens in successful immunotherapy of virally induced epithelial cancer. Science 356, 200–205. 10.1126/science.aak9510.

12. Lowery, F.J., Goff, S.L., Gasmi, B., Parkhurst, M.R., Ratnam, N.M., Halas, H.K., Shelton, T.E., Langhan, M.M., Bhasin, A., Dinerman, A.J., et al. (2025). Neoantigen-specific tumor-infiltrating lymphocytes in gastrointestinal cancers: a phase 2 trial. Nat Med 31, 1994–2003. 10.1038/s41591-025-03627-5.

13. Rizvi, N.A., Hellmann, M.D., Snyder, A., Kvistborg, P., Makarov, V., Havel, J.J., Lee, W., Yuan, J., Wong, P., Ho, T.S., et al. (2015). Cancer immunology. Mutational landscape determines sensitivity to PD-1 blockade in non-small cell lung cancer. Science 348, 124–128. 10.1126/science.aaa1348.

14. Samstein, R.M., Lee, C.-H., Shoushtari, A.N., Hellmann, M.D., Shen, R., Janjigian, Y.Y., Barron, D.A., Zehir, A., Jordan, E.J., Omuro, A., et al. (2019). Tumor mutational load predicts survival after immunotherapy across multiple cancer types. Nat Genet 51, 202–206. 10.1038/s41588-018-0312-8.

15. Gandara, D.R., Paul, S.M., Kowanetz, M., Schleifman, E., Zou, W., Li, Y., Rittmeyer, A., Fehrenbacher, L., Otto, G., Malboeuf, C., et al. (2018). Blood-based tumor mutational burden as a predictor of clinical benefit in non-small-cell lung cancer patients treated with atezolizumab. Nat Med 24, 1441–1448. 10.1038/s41591-018-0134-3.

16. Alban, T.J., Riaz, N., Parthasarathy, P., Makarov, V., Kendall, S., Yoo, S.-K., Shah, R., Weinhold, N., Srivastava, R., Ma, X., et al. (2024). Neoantigen immunogenicity landscapes and evolution of tumor ecosystems during immunotherapy with nivolumab. Nat Med, 1–14. 10.1038/s41591-024-03240-y.

17. Anagnostou, V., Smith, K.N., Forde, P.M., Niknafs, N., Bhattacharya, R., White, J., Zhang, T., Adleff, V., Phallen, J., Wali, N., et al. (2017). Evolution of Neoantigen Landscape during Immune Checkpoint Blockade in Non-Small Cell Lung Cancer. Cancer Discov 7, 264–276. 10.1158/2159-8290.CD-16-0828.

18. Riaz, N., Havel, J.J., Makarov, V., Desrichard, A., Urba, W.J., Sims, J.S., Hodi, F.S., Martín-Algarra, S., Mandal, R., Sharfman, W.H., et al. (2017). Tumor and Microenvironment Evolution during Immunotherapy with Nivolumab. Cell 171, 934–949.e16. 10.1016/j.cell.2017.09.028.

19. McGrail, D.J., Pilié, P.G., Rashid, N.U., Voorwerk, L., Slagter, M., Kok, M., Jonasch, E., Khasraw, M., Heimberger, A.B., Lim, B., et al. (2021). High tumor mutation burden fails to predict immune checkpoint blockade response across all cancer types. Ann Oncol 32, 661–672. 10.1016/j.annonc.2021.02.006.

20. Hu-Lieskovan, S., Lisberg, A., Zaretsky, J.M., Grogan, T.R., Rizvi, H., Wells, D.K., Carroll, J., Cummings, A., Madrigal, J., Jones, B., et al. (2019). Tumor Characteristics Associated with Benefit from Pembrolizumab in Advanced Non-Small Cell Lung Cancer. Clin Cancer Res 25, 5061–5068. 10.1158/1078-0432.CCR-18-4275.

21. Rozeman, E.A., Hoefsmit, E.P., Reijers, I.L.M., Saw, R.P.M., Versluis, J.M., Krijgsman, O., Dimitriadis, P., Sikorska, K., van de Wiel, B.A., Eriksson, H., et al. (2021). Survival and biomarker analyses from the OpACIN-neo and OpACIN neoadjuvant immunotherapy trials in stage III melanoma. Nat Med 27, 256–263. 10.1038/s41591-020-01211-7.

22. Hugo, W., Zaretsky, J.M., Sun, L., Song, C., Moreno, B.H., Hu-Lieskovan, S., Berent-Maoz, B., Pang, J., Chmielowski, B., Cherry, G., et al. (2016). Genomic and Transcriptomic Features of Response to Anti-PD-1 Therapy in Metastatic Melanoma. Cell 165, 35–44. 10.1016/j.cell.2016.02.065.

23. Mariathasan, S., Turley, S.J., Nickles, D., Castiglioni, A., Yuen, K., Wang, Y., Kadel, E.E., Koeppen, H., Astarita, J.L., Cubas, R., et al. (2018). TGFβ attenuates tumour response to PD-L1 blockade by contributing to exclusion of T cells. Nature 554, 544–548. 10.1038/nature25501.

24. Puig-Saus, C., Sennino, B., Peng, S., Wang, C.L., Pan, Z., Yuen, B., Purandare, B., An, D., Quach, B.B., Nguyen, D., et al. (2023). Neoantigen-targeted CD8+ T cell responses with PD-1 blockade therapy. Nature 615, 697–704. 10.1038/s41586-023-05787-1.

25. Caushi, J.X., Zhang, J., Ji, Z., Vaghasia, A., Zhang, B., Hsiue, E.H.-C., Mog, B.J., Hou, W., Justesen, S., Blosser, R., et al. (2021). Transcriptional programs of neoantigen-specific TIL in anti-PD-1-treated lung cancers. Nature 596, 126–132. 10.1038/s41586-021-03752-4.

26. Balança, C.-C., Scarlata, C.-M., Michelas, M., Devaud, C., Sarradin, V., Franchet, C., Martinez Gomez, C., Gomez-Roca, C., Tosolini, M., Heaugwane, D., et al. (2020). Dual Relief of T-lymphocyte Proliferation and Effector Function Underlies Response to PD-1 Blockade in Epithelial Malignancies. Cancer Immunology Research 8, 869–882. 10.1158/2326-6066.CIR-19-0855.

27. Thommen, D.S., Koelzer, V.H., Herzig, P., Roller, A., Trefny, M., Dimeloe, S., Kiialainen, A., Hanhart, J., Schill, C., Hess, C., et al. (2018). A transcriptionally and functionally distinct PD-1+ CD8+ T cell pool with predictive potential in non-small-cell lung cancer treated with PD-1 blockade. Nat. Med. 24, 994–1004. 10.1038/s41591-018-0057-z.

28. Metoikidou, C., Karnaukhov, V., Boeckx, B., Timperi, E., Bonté, P.-E., Wang, L., Espenel, M., Albaud, B., Loirat, D., Wang, X., et al. (2025). Continuous replenishment of the dysfunctional CD8 T cell axis is associated with response to chemoimmunotherapy in advanced breast cancer. Cell Reports Medicine 6, 101973. 10.1016/j.xcrm.2025.101973.

29. Kaptein, P., Slingerland, N., Van Der Leun, A.M., Runderkamp, E., Wagensveld, R.A., Michael Chin, S., Mors, J.R., Machuca-Ostos, M., Reissig, T.M., Moynihan, K.D., et al. (2025). Reinvigoration of translational activity in dysfunctional T cells initiates the early intratumoral response to PD-1 blockade. Preprint, 10.1101/2025.09.24.676875.

30. Sade-Feldman, M., Yizhak, K., Bjorgaard, S.L., Ray, J.P., de Boer, C.G., Jenkins, R.W., Lieb, D.J., Chen, J.H., Frederick, D.T., Barzily-Rokni, M., et al. (2018). Defining T Cell States Associated with Response to Checkpoint Immunotherapy in Melanoma. Cell 175, 998–1013.e20. 10.1016/j.cell.2018.10.038.

31. Balança, C.-C., Salvioni, A., Scarlata, C.-M., Michelas, M., Martinez-Gomez, C., Gomez-Roca, C., Sarradin, V., Tosolini, M., Valle, C., Pont, F., et al. (2021). PD-1 blockade restores helper activity of tumor-infiltrating, exhausted PD-1hiCD39+ CD4 T cells. JCI Insight 6, e142513. 10.1172/jci.insight.142513.

32. Yossef, R., Krishna, S., Sindiri, S., Lowery, F.J., Copeland, A.R., Gartner, J.J., Parkhurst, M.R., Parikh, N.B., Hitscherich, K.J., Levi, S.T., et al. (2023). Phenotypic signatures of circulating neoantigen-reactive CD8+ T cells in patients with metastatic cancers. Cancer Cell 41, 2154–2165.e5. 10.1016/j.ccell.2023.11.005.

33. Ellrott, K., Bailey, M.H., Saksena, G., Covington, K.R., Kandoth, C., Stewart, C., Hess, J., Ma, S., Chiotti, K.E., McLellan, M., et al. (2018). Scalable Open Science Approach for Mutation Calling of Tumor Exomes Using Multiple Genomic Pipelines. Cell Syst 6, 271–281.e7. 10.1016/j.cels.2018.03.002.

34. Wells, D.K., van Buuren, M.M., Dang, K.K., Hubbard-Lucey, V.M., Sheehan, K.C.F., Campbell, K.M., Lamb, A., Ward, J.P., Sidney, J., Blazquez, A.B., et al. (2020). Key Parameters of Tumor Epitope Immunogenicity Revealed Through a Consortium Approach Improve Neoantigen Prediction. Cell 183, 818–834.e13. 10.1016/j.cell.2020.09.015.

35. Kawaguchi, S., Higasa, K., Shimizu, M., Yamada, R., and Matsuda, F. (2017). HLA-HD: An accurate HLA typing algorithm for next-generation sequencing data. Hum Mutat 38, 788–797. 10.1002/humu.23230.

36. Thrift, W.J., Lounsbury, N.W., Broadwell, Q., Heidersbach, A., Freund, E., Abdolazimi, Y., Phung, Q.T., Chen, J., Capietto, A.-H., Tong, A.-J., et al. (2024). Towards designing improved cancer immunotherapy targets with a peptide-MHC-I presentation model, HLApollo. Nat Commun 15, 10752. 10.1038/s41467-024-54887-7.

37. Reynisson, B., Barra, C., Kaabinejadian, S., Hildebrand, W.H., Peters, B., and Nielsen, M. (2020). Improved Prediction of MHC II Antigen Presentation through Integration and Motif Deconvolution of Mass Spectrometry MHC Eluted Ligand Data. J Proteome Res 19, 2304–2315. 10.1021/acs.jproteome.9b00874.

38. Ravi, A., Hellmann, M.D., Arniella, M.B., Holton, M., Freeman, S.S., Naranbhai, V., Stewart, C., Leshchiner, I., Kim, J., Akiyama, Y., et al. (2023). Genomic and transcriptomic analysis of checkpoint blockade response in advanced non-small cell lung cancer. Nat Genet 55, 807–819. 10.1038/s41588-023-01355-5.

39. Eberhardt, C.S., Kissick, H.T., Patel, M.R., Cardenas, M.A., Prokhnevska, N., Obeng, R.C., Nasti, T.H., Griffith, C.C., Im, S.J., Wang, X., et al. (2021). Functional HPV-specific PD-1+ stem-like CD8 T cells in head and neck cancer. Nature 597, 279–284. 10.1038/s41586-021-03862-z.

40. Schumacher, T.N., and Schreiber, R.D. (2015). Neoantigens in cancer immunotherapy. Science 348, 69–74. 10.1126/science.aaa4971.

41. Hudson, W.H., Gensheimer, J., Hashimoto, M., Wieland, A., Valanparambil, R.M., Li, P., Lin, J.-X., Konieczny, B.T., Im, S.J., Freeman, G.J., et al. (2019). Proliferating Transitory T Cells with an Effector-like Transcriptional Signature Emerge from PD-1+ Stem-like CD8+ T Cells during Chronic Infection. Immunity 51, 1043–1058.e4. 10.1016/j.immuni.2019.11.002.

42. Hu, W., Shawn Hu, S., Zhu, S., Peng, W., Badovinac, V.P., Zang, C., Zhao, X., and Xue, H.-H. (2025). Hdac1 as an early determinant of intermediate-exhausted CD8+ T cell fate in chronic viral infection. Proc Natl Acad Sci U S A 122, e2502256122. 10.1073/pnas.2502256122.

43. Im, S.J., Hashimoto, M., Gerner, M.Y., Lee, J., Kissick, H.T., Burger, M.C., Shan, Q., Hale, J.S., Lee, J., Nasti, T.H., et al. (2016). Defining CD8+ T cells that provide the proliferative burst after PD-1 therapy. Nature 537, 417–421. 10.1038/nature19330.

44. Weulersse, M., Asrir, A., Pichler, A.C., Lemaitre, L., Braun, M., Carrié, N., Joubert, M.-V., Le Moine, M., Do Souto, L., Gaud, G., et al. (2020). Eomes-Dependent Loss of the Co-activating Receptor CD226 Restrains CD8+ T Cell Anti-tumor Functions and Limits the Efficacy of Cancer Immunotherapy. Immunity 53, 824–839.e10. 10.1016/j.immuni.2020.09.006.

45. Hui, E., Cheung, J., Zhu, J., Su, X., Taylor, M.J., Wallweber, H.A., Sasmal, D.K., Huang, J., Kim, J.M., Mellman, I., et al. (2017). T cell costimulatory receptor CD28 is a primary target for PD-1-mediated inhibition. Science 355, 1428–1433. 10.1126/science.aaf1292.

46. Wang, G., Yoon, D., Nandi, A., Patel, K., Azar, T., Kim, J., Han, N.A., Nickie, A., Park, S., Wang, K., et al. (2026). Antigen-specific profiling identifies T-bet+ melanoma-specific CD8+ T cells associated with response to neoadjuvant PD-1 blockade. Cancer Cell 44, 221–234.e5. 10.1016/j.ccell.2025.12.004.

47. Gros, A., Parkhurst, M.R., Tran, E., Pasetto, A., Robbins, P.F., Ilyas, S., Prickett, T.D., Gartner, J.J., Crystal, J.S., Roberts, I.M., et al. (2016). Prospective identification of neoantigen-specific lymphocytes in the peripheral blood of melanoma patients. Nat Med 22, 433–438. 10.1038/nm.4051.

48. Siddiqui, I., Schaeuble, K., Chennupati, V., Fuertes Marraco, S.A., Calderon-Copete, S., Pais Ferreira, D., Carmona, S.J., Scarpellino, L., Gfeller, D., Pradervand, S., et al. (2019). Intratumoral Tcf1+PD-1+CD8+ T Cells with Stem-like Properties Promote Tumor Control in Response to Vaccination and Checkpoint Blockade Immunotherapy. Immunity 50, 195–211.e10. 10.1016/j.immuni.2018.12.021.

49. McLane, L.M., Abdel-Hakeem, M.S., and Wherry, E.J. (2019). CD8 T Cell Exhaustion During Chronic Viral Infection and Cancer. Annu. Rev. Immunol. 37, 457–495. 10.1146/annurev-immunol-041015-055318.

50. Kamphorst, A.O., Wieland, A., Nasti, T., Yang, S., Zhang, R., Barber, D.L., Konieczny, B.T., Daugherty, C.Z., Koenig, L., Yu, K., et al. (2017). Rescue of exhausted CD8 T cells by PD-1-targeted therapies is CD28-dependent. Science 355, 1423–1427. 10.1126/science.aaf0683.

51. Banta, K.L., Xu, X., Chitre, A.S., Au-Yeung, A., Takahashi, C., O’Gorman, W.E., Wu, T.D., Mittman, S., Cubas, R., Comps-Agrar, L., et al. (2022). Mechanistic convergence of the TIGIT and PD-1 inhibitory pathways necessitates co-blockade to optimize anti-tumor CD8+ T cell responses. Immunity 55, 512–526.e9. 10.1016/j.immuni.2022.02.005.

52. Huang, Q., Wu, X., Wang, Z., Chen, X., Wang, L., Lu, Y., Xiong, D., Liu, Q., Tian, Y., Lin, H., et al. (2022). The primordial differentiation of tumor-specific memory CD8+ T cells as bona fide responders to PD-1/PD-L1 blockade in draining lymph nodes. Cell 185, 4049–4066.e25. 10.1016/j.cell.2022.09.020.

53. Honigsberg, R., Cruz, T., Yoffe, L., Tang, M.S., Dicle, O., Markowitz, G., Michael, M., Singh, A., Altorki, N.K., Elemento, O., et al. (2025). Tumor-specific draining lymph node CD8 T cells orchestrate an anti-tumor response to neoadjuvant PD-1 immune checkpoint blockade. bioRxiv, 2025.04.27.650862. 10.1101/2025.04.27.650862.

54. Mikucki, M.E., Fisher, D.T., Matsuzaki, J., Skitzki, J.J., Gaulin, N.B., Muhitch, J.B., Ku, A.W., Frelinger, J.G., Odunsi, K., Gajewski, T.F., et al. (2015). Non-redundant requirement for CXCR3 signalling during tumoricidal T-cell trafficking across tumour vascular checkpoints. Nat Commun 6, 7458. 10.1038/ncomms8458.

55. Voskoboinik, I., Whisstock, J.C., and Trapani, J.A. (2015). Perforin and granzymes: function, dysfunction and human pathology. Nat Rev Immunol 15, 388–400. 10.1038/nri3839.

56. Rojas, L.A., Sethna, Z., Soares, K.C., Olcese, C., Pang, N., Patterson, E., Lihm, J., Ceglia, N., Guasp, P., Chu, A., et al. (2023). Personalized RNA neoantigen vaccines stimulate T cells in pancreatic cancer. Nature 618, 144–150. 10.1038/s41586-023-06063-y.

57. Weber, J.S., Carlino, M.S., Khattak, A., Meniawy, T., Ansstas, G., Taylor, M.H., Kim, K.B., McKean, M., Long, G.V., Sullivan, R.J., et al. (2024). Individualised neoantigen therapy mRNA-4157 (V940) plus pembrolizumab versus pembrolizumab monotherapy in resected melanoma (KEYNOTE-942): a randomised, phase 2b study. Lancet 403, 632–644. 10.1016/S0140-6736(23)02268-7.

58. Blass, E., Keskin, D.B., Tu, C.R., Forman, C., Vanasse, A., Sax, H.E., Shim, B., Chea, V., Kim, N., Carulli, I., et al. (2025). A multi-adjuvant personal neoantigen vaccine generates potent immunity in melanoma. Cell 188, 5125–5141.e27. 10.1016/j.cell.2025.06.019.

59. Sahin, U., Derhovanessian, E., Miller, M., Kloke, B.-P., Simon, P., Löwer, M., Bukur, V., Tadmor, A.D., Luxemburger, U., Schrörs, B., et al. (2017). Personalized RNA mutanome vaccines mobilize poly-specific therapeutic immunity against cancer. Nature 547, 222–226. 10.1038/nature23003.

60. Ott, P.A., Hu, Z., Keskin, D.B., Shukla, S.A., Sun, J., Bozym, D.J., Zhang, W., Luoma, A., Giobbie-Hurder, A., Peter, L., et al. (2017). An immunogenic personal neoantigen vaccine for patients with melanoma. Nature 547, 217–221. 10.1038/nature22991.

61. Lang, F., Schrörs, B., Löwer, M., Türeci, Ö., and Sahin, U. (2022). Identification of neoantigens for individualized therapeutic cancer vaccines. Nat Rev Drug Discov 21, 261–282. 10.1038/s41573-021-00387-y.

62. Borst, J., Ahrends, T., Bąbała, N., Melief, C.J.M., and Kastenmüller, W. (2018). CD4+ T cell help in cancer immunology and immunotherapy. Nat. Rev. Immunol. 18, 635–647. 10.1038/s41577-018-0044-0.

63. Alspach, E., Lussier, D.M., Miceli, A.P., Kizhvatov, I., DuPage, M., Luoma, A.M., Meng, W., Lichti, C.F., Esaulova, E., Vomund, A.N., et al. (2019). MHC-II neoantigens shape tumour immunity and response to immunotherapy. Nature 574, 696–701. 10.1038/s41586-019-1671-8.

64. Tran, E., Turcotte, S., Gros, A., Robbins, P.F., Lu, Y.-C., Dudley, M.E., Wunderlich, J.R., Somerville, R.P., Hogan, K., Hinrichs, C.S., et al. (2014). Cancer Immunotherapy Based on Mutation-Specific CD4+ T Cells in a Patient with Epithelial Cancer. Science 344, 641–645. 10.1126/science.1251102.

65. Gfeller, D., Schmidt, J., Croce, G., Guillaume, P., Bobisse, S., Genolet, R., Queiroz, L., Cesbron, J., Racle, J., and Harari, A. (2023). Improved predictions of antigen presentation and TCR recognition with MixMHCpred2.2 and PRIME2.0 reveal potent SARS-CoV-2 CD8+ T-cell epitopes. Cell Syst 14, 72–83.e5. 10.1016/j.cels.2022.12.002.

66. Li, H., and Durbin, R. (2009). Fast and accurate short read alignment with Burrows–Wheeler transform. Bioinformatics 25, 1754–1760. 10.1093/bioinformatics/btp324.

67. Cibulskis, K., Lawrence, M.S., Carter, S.L., Sivachenko, A., Jaffe, D., Sougnez, C., Gabriel, S., Meyerson, M., Lander, E.S., and Getz, G. (2013). Sensitive detection of somatic point mutations in impure and heterogeneous cancer samples. Nat Biotechnol 31, 213–219. 10.1038/nbt.2514.

68. Saunders, C.T., Wong, W.S.W., Swamy, S., Becq, J., Murray, L.J., and Cheetham, R.K. (2012). Strelka: accurate somatic small-variant calling from sequenced tumor–normal sample pairs. Bioinformatics 28, 1811–1817. 10.1093/bioinformatics/bts271.

69. Wilm, A., Aw, P.P.K., Bertrand, D., Yeo, G.H.T., Ong, S.H., Wong, C.H., Khor, C.C., Petric, R., Hibberd, M.L., and Nagarajan, N. (2012). LoFreq: a sequence-quality aware, ultra-sensitive variant caller for uncovering cell-population heterogeneity from high-throughput sequencing datasets. Nucleic Acids Research 40, 11189–11201. 10.1093/nar/gks918.

70. Alexandrov, L.B., Kim, J., Haradhvala, N.J., Huang, M.N., Tian Ng, A.W., Wu, Y., Boot, A., Covington, K.R., Gordenin, D.A., Bergstrom, E.N., et al. (2020). The repertoire of mutational signatures in human cancer. Nature 578, 94–101. 10.1038/s41586-020-1943-3.

71. Dobin, A., Davis, C.A., Schlesinger, F., Drenkow, J., Zaleski, C., Jha, S., Batut, P., Chaisson, M., and Gingeras, T.R. (2013). STAR: ultrafast universal RNA-seq aligner. Bioinformatics 29, 15–21. 10.1093/bioinformatics/bts635.

72. Anders, S., Pyl, P.T., and Huber, W. (2015). HTSeq—a Python framework to work with high-throughput sequencing data. Bioinformatics 31, 166–169. 10.1093/bioinformatics/btu638.

73. Genentech (2025). Genentech/rnacounts: v1. Version v1 (Zenodo). 10.5281/ZENODO.17060247.

74. Gu, Z., Eils, R., and Schlesner, M. (2016). Complex heatmaps reveal patterns and correlations in multidimensional genomic data. Bioinformatics 32, 2847–2849. 10.1093/bioinformatics/btw313.

75. Hänzelmann, S., Castelo, R., and Guinney, J. (2013). GSVA: gene set variation analysis for microarray and RNA-Seq data. BMC Bioinformatics 14, 7. 10.1186/1471-2105-14-7.

76. Newman, A.M., Liu, C.L., Green, M.R., Gentles, A.J., Feng, W., Xu, Y., Hoang, C.D., Diehn, M., and Alizadeh, A.A. (2015). Robust enumeration of cell subsets from tissue expression profiles. Nat Methods 12, 453–457. 10.1038/nmeth.3337.

77. Ayers, M., Lunceford, J., Nebozhyn, M., Murphy, E., Loboda, A., Kaufman, D.R., Albright, A., Cheng, J.D., Kang, S.P., Shankaran, V., et al. (2017). IFN-γ–related mRNA profile predicts clinical response to PD-1 blockade. Journal of Clinical Investigation 127, 2930–2940. 10.1172/JCI91190.

78. Lam, K.C., Araya, R.E., Huang, A., Chen, Q., Di Modica, M., Rodrigues, R.R., Lopès, A., Johnson, S.B., Schwarz, B., Bohrnsen, E., et al. (2021). Microbiota triggers STING-type I IFN-dependent monocyte reprogramming of the tumor microenvironment. Cell 184, 5338–5356.e21. 10.1016/j.cell.2021.09.019.

79. Van Loo, P., Nordgard, S.H., Lingjærde, O.C., Russnes, H.G., Rye, I.H., Sun, W., Weigman, V.J., Marynen, P., Zetterberg, A., Naume, B., et al. (2010). Allele-specific copy number analysis of tumors. Proc. Natl. Acad. Sci. U.S.A. 107, 16910–16915. 10.1073/pnas.1009843107.

80. Nik-Zainal, S., Van Loo, P., Wedge, D.C., Alexandrov, L.B., Greenman, C.D., Lau, K.W., Raine, K., Jones, D., Marshall, J., Ramakrishna, M., et al. (2012). The Life History of 21 Breast Cancers. Cell 149, 994–1007. 10.1016/j.cell.2012.04.023.

81. Dentro, S.C., Wedge, D.C., and Van Loo, P. (2017). Principles of Reconstructing the Subclonal Architecture of Cancers. Cold Spring Harb Perspect Med 7, a026625. 10.1101/cshperspect.a026625.

82. McLaren, W., Gil, L., Hunt, S.E., Riat, H.S., Ritchie, G.R.S., Thormann, A., Flicek, P., and Cunningham, F. (2016). The Ensembl Variant Effect Predictor. Genome Biol 17, 122. 10.1186/s13059-016-0974-4.

83. Reynisson, B., Alvarez, B., Paul, S., Peters, B., and Nielsen, M. (2020). NetMHCpan-4.1 and NetMHCIIpan-4.0: improved predictions of MHC antigen presentation by concurrent motif deconvolution and integration of MS MHC eluted ligand data. Nucleic Acids Research 48, W449–W454. 10.1093/nar/gkaa379.

84. Tadros, D.M., Racle, J., and Gfeller, D. (2025). Predicting MHC-I ligands across alleles and species: how far can we go? Genome Med 17, 25. 10.1186/s13073-025-01450-8.

85. Harndahl, M., Rasmussen, M., Roder, G., Dalgaard Pedersen, I., Sørensen, M., Nielsen, M., and Buus, S. (2012). Peptide- MHC class I stability is a better predictor than peptide affinity of CTL immunogenicity. Eur J Immunol 42, 1405–1416. 10.1002/eji.201141774.

86. Nielsen, M., Lundegaard, C., Lund, O., and Keşmir, C. (2005). The role of the proteasome in generating cytotoxic T-cell epitopes: insights obtained from improved predictions of proteasomal cleavage. Immunogenetics 57, 33–41. 10.1007/s00251-005-0781-7.

87. Lu, Y.-C., Yao, X., Crystal, J.S., Li, Y.F., El-Gamil, M., Gross, C., Davis, L., Dudley, M.E., Yang, J.C., Samuels, Y., et al. (2014). Efficient Identification of Mutated Cancer Antigens Recognized by T Cells Associated with Durable Tumor Regressions. Clinical Cancer Research 20, 3401–3410. 10.1158/1078-0432.CCR-14-0433.

88. Gurung, H.R., Heidersbach, A.J., Darwish, M., Chan, P.P.F., Li, J., Beresini, M., Zill, O.A., Wallace, A., Tong, A.-J., Hascall, D., et al. (2024). Systematic discovery of neoepitope-HLA pairs for neoantigens shared among patients and tumor types. Nat Biotechnol 42, 1107–1117. 10.1038/s41587-023-01945-y.

89. Darwish, M., Wichner, S., Li, J., Chang, J.C., Tam, C., Franke, Y., Li, H., Chan, P., and Blanchette, C. (2021). High-throughput identification of conditional MHCI ligands and scaled-up production of conditional MHCI complexes. Protein Sci 30, 1169–1183. 10.1002/pro.4082.

